# Comparative anatomy of the giraffe distal limb

**DOI:** 10.64898/2026.04.18.719342

**Authors:** Ray Wilhite, Diana Miller, Kelsey Newman, Stephanie Fennessy, Julian Fennessy, Michael Butler Brown, Amy Schilz

## Abstract

Giraffe in human care are known to experience significant clinical issues related to their feet. To characterize normal foot anatomy, we analyzed six sets of front and hind feet from wild Angolan giraffe and one calf in human care. We used computed tomography, three-dimensional reconstruction, sagittal sections, and gross dissection to acquire as much gross anatomical detail as possible. Significant anatomical findings include the deep digital flexor tendon that is very gracile as it crosses the fetlock and proximal phalanges but widens significantly before inserting onto the entire palmar/plantar surface of the distal phalanx. Significant subcutaneous abaxial veins were noted at the level of the fetlock in both front and hind feet. The digital cushion was found to be a complex structure consisting of two distinct regions, one underneath the distal phalanx, characterized by multiple transversely oriented small adipose compartments separated by dense connective tissue septa, and a significantly larger portion within the heel bulb, consisting of two sagittally oriented fat bodies encased in a dense connective tissue capsule and divided by a thick septum. The proportion of adipose tissue volume in the heel bulbs compared to the distal phalanx decreases with age. The thickness of the sole was found to be much greater than that of the wall and the sole appears to be the major weight supporting structure of the foot. In particular, the heel is greatly expanded in giraffe relative to other ruminants and was found to consist of softer material than the rest of the sole. Data presented here provide an overview of normal giraffe foot anatomy, which can be compared with data from giraffe in human care to better understand, guide treatment and prevention of abnormal anatomical conditions.

## Introduction

Approximately 1,000 giraffe (*Giraffa spp.*) are believed to be in human care in the United States of America, and over 2,000 giraffe worldwide [1]. More specifically, the studbooks indicate giraffe are distributed across 465 institutions, with 804 in USDA-registered facilities in 2022 [2], 573 of which were held in facilities accredited by the Association of Zoos and Aquariums (AZA) (pers. comm. Amy Schilz, February 2026), and 296 in facilities accredited by the Zoological Association of America (ZAA) (pers. comm. Tiffany Soechting, January 2026), [3, 4]. Giraffe in human care often experience significant hoof-related issues leading to health concerns, including overgrowth and lameness [1, 5]. With these chronic hoof-related issues, the weight distribution can be altered and lead to secondary pathologies and irregular hoof shape [5]. Clinical conditions associated with their feet present significant challenges to long-term maintenance under human care [5, 6], and understanding presentations of distal limb anatomy, under both in situ and ex situ conditions, is critical for informing effective hoof care management programs.

While giraffe voluntary training hoof care programs have become a standard of care [7], trimming practices that address giraffe hoof pathologies are currently based on domestic ruminants and horses [8]. The knowledge gap between domestic and exotic animal hoof care has begun to narrow with increased data collection, particularly studies characterizing the giraffe hoof capsule and its radiographic anatomy [5, 7, 9]. However, with the exception of a single dissertation [10], few studies have evaluated the soft tissue anatomy of the distal limb. These soft tissues are integral in understanding the form and function of the hoof and, in turn, for informing proper hoof health management.

Dadone et al. [11] characterized the macroscopic and microscopic anatomy of hybrid southern giraffe (*G. giraffa)* hoof capsule, dermis and digital cushion. While not related directly to giraffe, a study of the bovine digital cushion by Raber et al. [12] showed a direct correlation between increased loading of the foot in pregnant dairy cattle (*Bos taurus*) and decreasing fat content of the digital cushion. Here, we present a further detailed interpretation of the digital cushion. We hypothesize a similar correlation may be present in the heel bulbs of healthy maturing giraffe as weight increases with age. To facilitate this comparison with cattle, we also examined differences between their distal limb anatomy. This comparison is especially useful considering bovine and equine feet are frequently used as analogs for giraffe when assessing footcare. Raber et al. [12] further noted that in cattle, the adipose bodies within the claws were asymmetrically distributed, with the front lateral claw having more adipose tissue than the front medial claw and the rear medial claw having more adipose tissue than the rear lateral claw. While this distribution was expected to be similar in giraffe, Muirhead et al. [13] examined the biomechanics of sound feet in zoo giraffe and highlighted that the lateral claw was the main weight-bearing claw most of the time. Studies in cattle have shown that the lateral claw in the hind limb bears most of the weight, but in the front limb, weight seems to be more evenly distributed, with the medial claw being slightly favored [14].

A limitation of the Muirhead et al. [13] study was the reliance on zoo-housed giraffe and the fact that the ground surfaces of the feet could not be examined to confirm there were no indications of hoof overgrowth. Our study examines the trends observed in wild giraffe foot asymmetries, as they should best correspond to normal giraffe foot anatomy. We also present a proposed standard measurement scheme for the hoof capsule and a detailed description of the anatomy of the giraffe distal limb based on multiple imaging modalities. We used wild Angolan giraffe specimens (*G. giraffa angolensis*) as well as a single 2.5 month old northern giraffe (*G. camelopardalis*) human care specimen to establish a baseline of normal anatomy with which to compare animals with clinical foot issues in the future. Our work provides novel insights into the nature of the digital cushion and redefines the structure in giraffe. We also define a unique region of the caudal digital cushion as the heel bulb and give a detailed description of its anatomy.

## Materials and Methods

One northern giraffe in human care and six wild Angolan giraffe (*G. giraffa angolensis*) were used for this study. The single human care specimen was a 2.5 month old female northern giraffe (*G. camelopardalis*) with no history of foot problems.We hypothesized that her feet should be virtually identical to those of similar aged wild giraffe. This specimen was important for establishing a baseline to which the older wild specimens could be compared. Of the wild specimens, five were subadult males sustainably harvested as part of a conservation management program at Okonjati Game Reserve in central Namibia, and another from a wild female which died from unknown causes unrelated to foot trauma at Etosha Heights Private Reserve, north-central Namibia. Distal front and rear limbs were removed approximately 15 cm above the fetlock.

To better understand the anatomy of the foot, both computed tomography (CT) and anatomical dissection data were collected. CT data were collected from all four limbs for each individual. Wild giraffe feet were scanned in Windhoek, Namibia at 0.6 mm slice intervals on a Siemens Somaton Definition 64 slice scanner, Serial 67278 (Medical Imaging Namibia), while human care feet were scanned at 0.625 mm slice intervals on a GE Revolution EVO 128 CT (Auburn University College of Veterinary Medicine). CT data were processed using Materialize Mimics 25. For all scans, the distal phalanx of each digit was auto segmented and rendered as a solid model and the heel bulbs were manually segmented using the “Multiple Slice Edit” tool in Mimics (Materialize Mimics version 25). Manual segmentation was based on visual separation of the adipose-containing portion of the heel bulb from the dense connective tissue capsule (Fig 1a). The heel bulbs were then rendered as solid models for comparison with the distal phalanx (Fig 2). For select scans, extensors and flexors were also segmented using the “Multiple Slice Edit” tool in order to visualize the geometry of the individual tendons (Fig 3).

**Figure 1.**
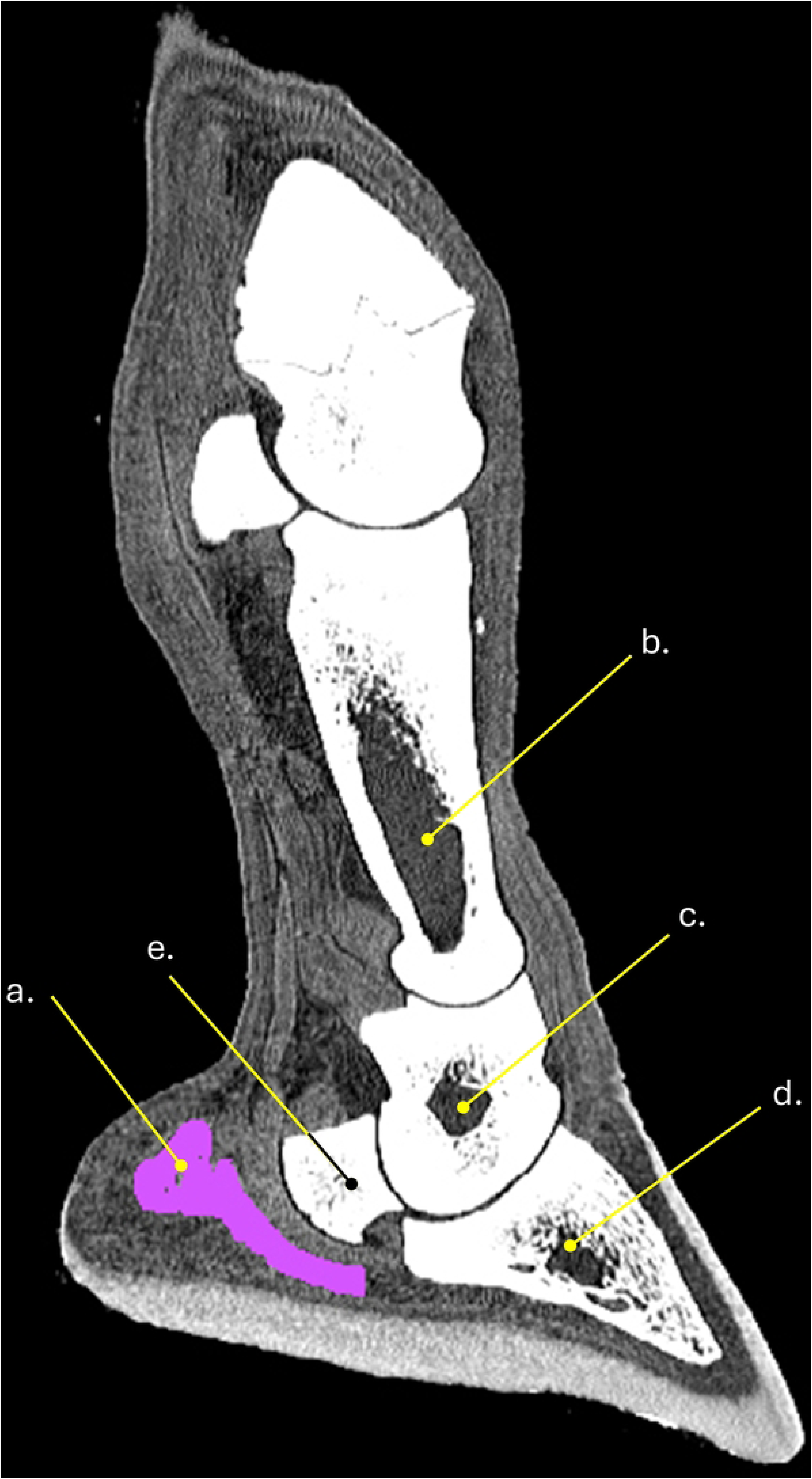
CT mid-sagittal section through the medial claw of the right thoracic limb of a wild Angolan giraffe. a- adipose component of heel bulb. b- medullary cavity of proximal phalanx. c- medullary cavity of middle phalanx. d- medullary cavity of distal phalanx.

**Figure 2.**
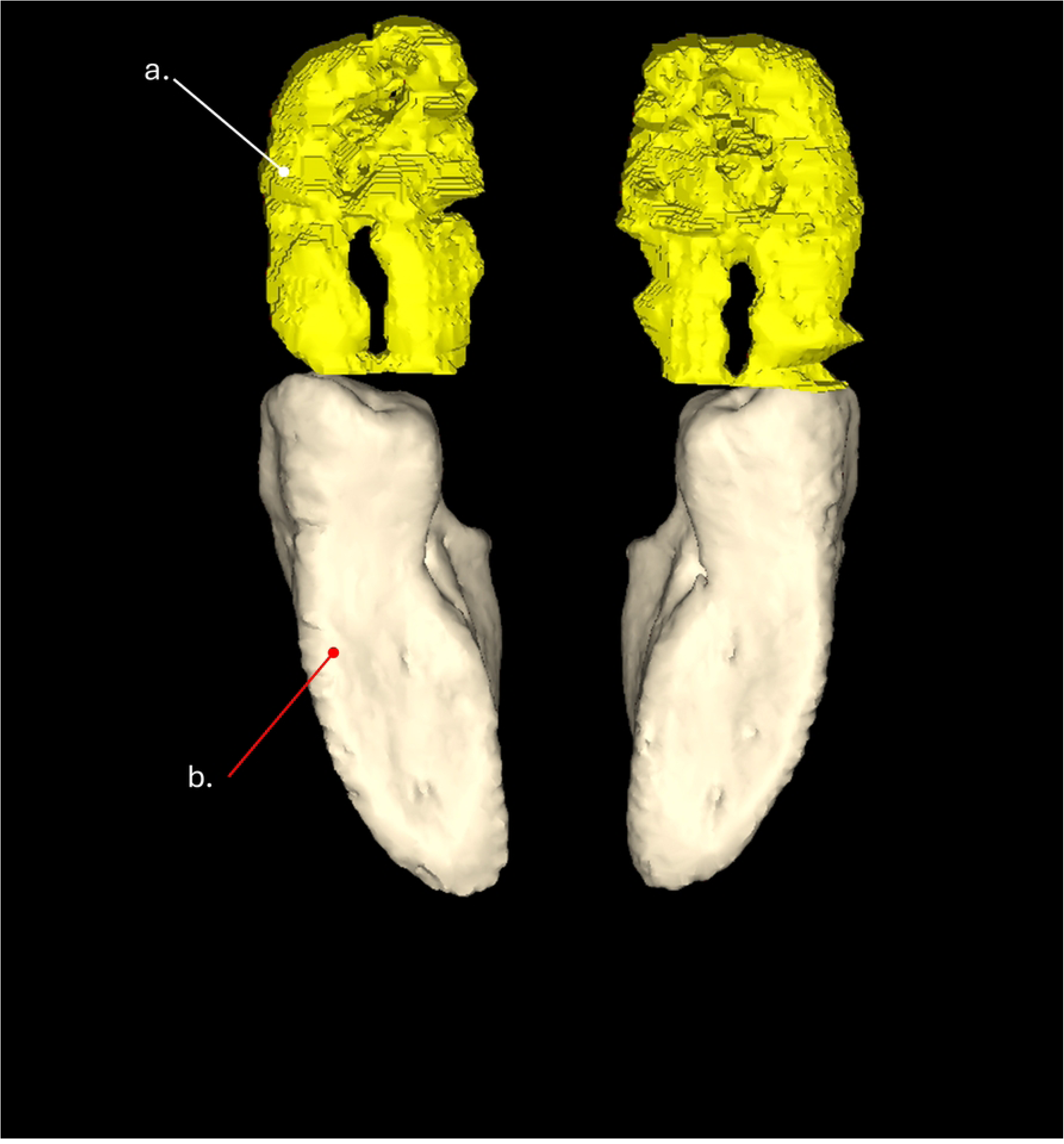
3-D reconstruction of a right thoracic distal limb in a distal view of a wild Angolan giraffe. a- adipose tissue of the heel bulb. b- distal phalanx.

**Figure 3.**
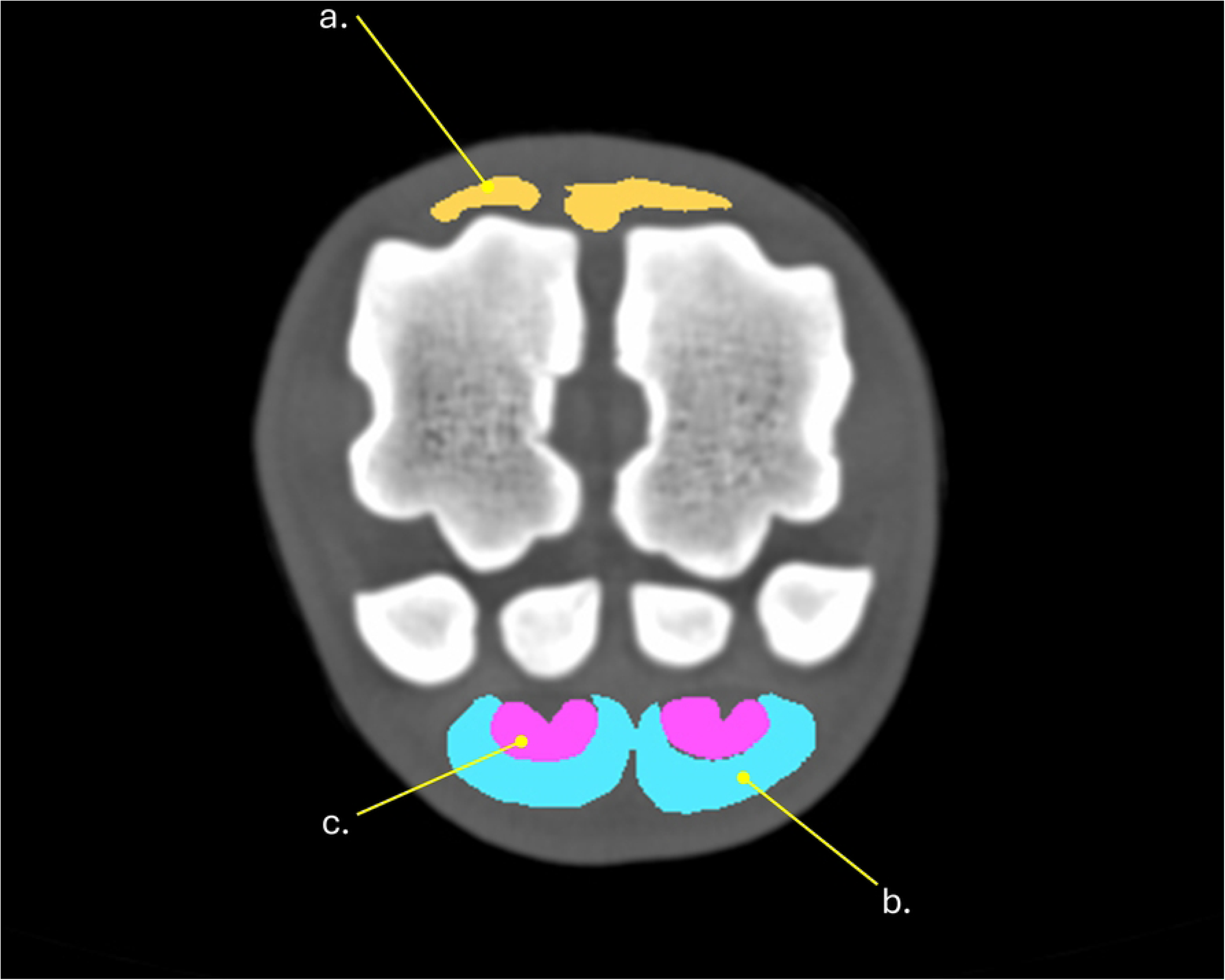
CT cross-section through the mid-fetlock joint at the level of the proximal sesamoids in a right pelvic limb of a wild Angolan giraffe. a- digital extensor ligaments. b-superficial digital flexor tendons. c-deep digital flexor tendons.

Before dissection or sectioning, each foot was photographed in dorsal, palmar/plantar, medial, lateral and distal views (Fig 4). Five measurements were made for each hoof capsule consisting of claw length, heel width, median width, toe height and toe width (Fig 5). Total width of the combined hoof capsules was also recorded. Left limbs were sectioned using a band saw. For each sectioned foot, cuts were made in a sagittal plane through the interdigital space as well as through the center of each digit for a total of three cuts per foot (Fig 6). Both sides of each cut were photographed and documented. Photos of the human care specimen were taken with the camera locked on a copy stand to get perpendicular photos, and the camera held as close as possible to perpendicular for photos of wild specimens.

**Figure 4.**
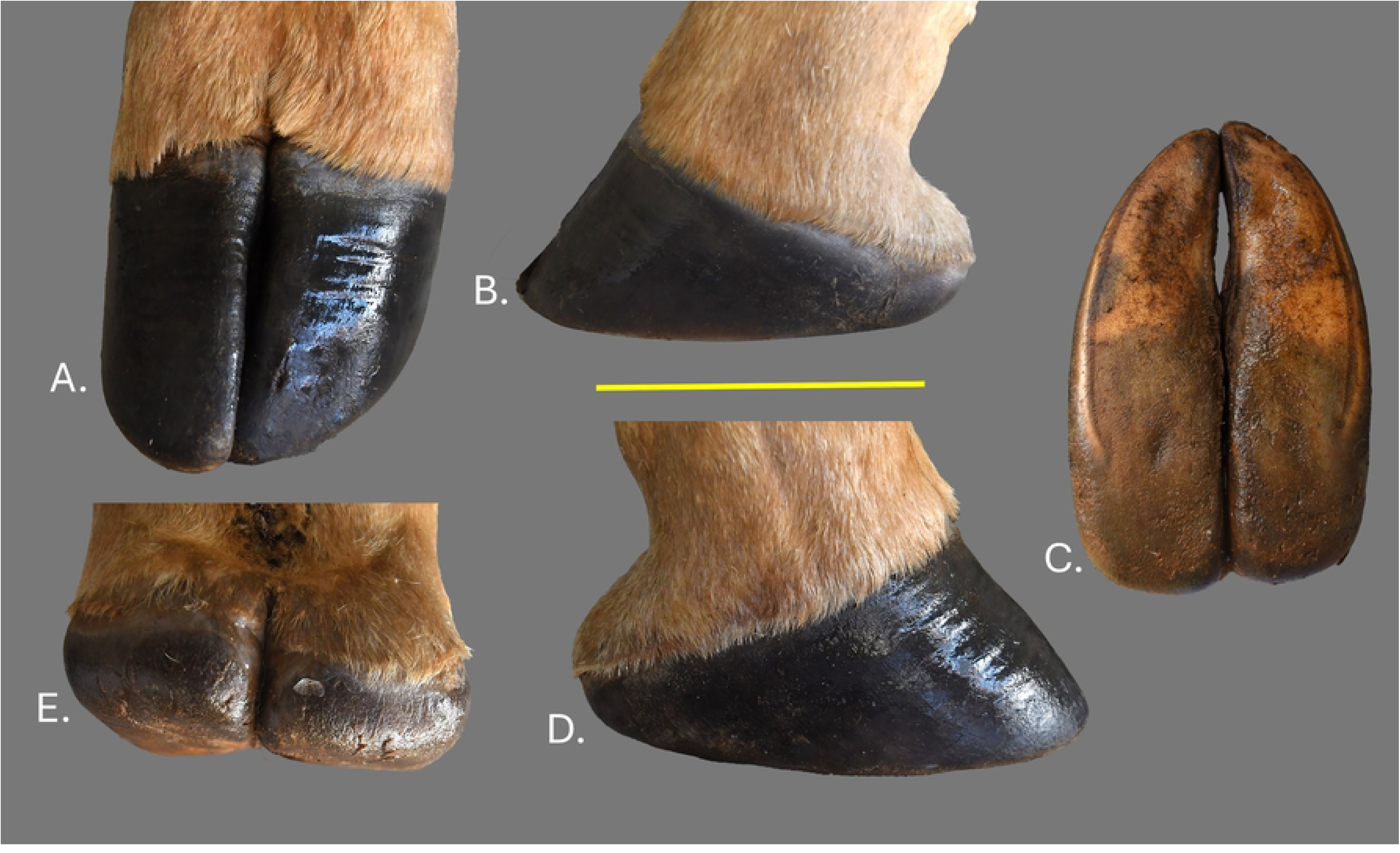
Right thoracic foot of a wild Angolan giraffe. (A) Dorsal view. (B) Medial view. (C) Distal view. (D) Lateral view. (E) Palmar view. Scale bar equals 10cm.

**Figure 5.**
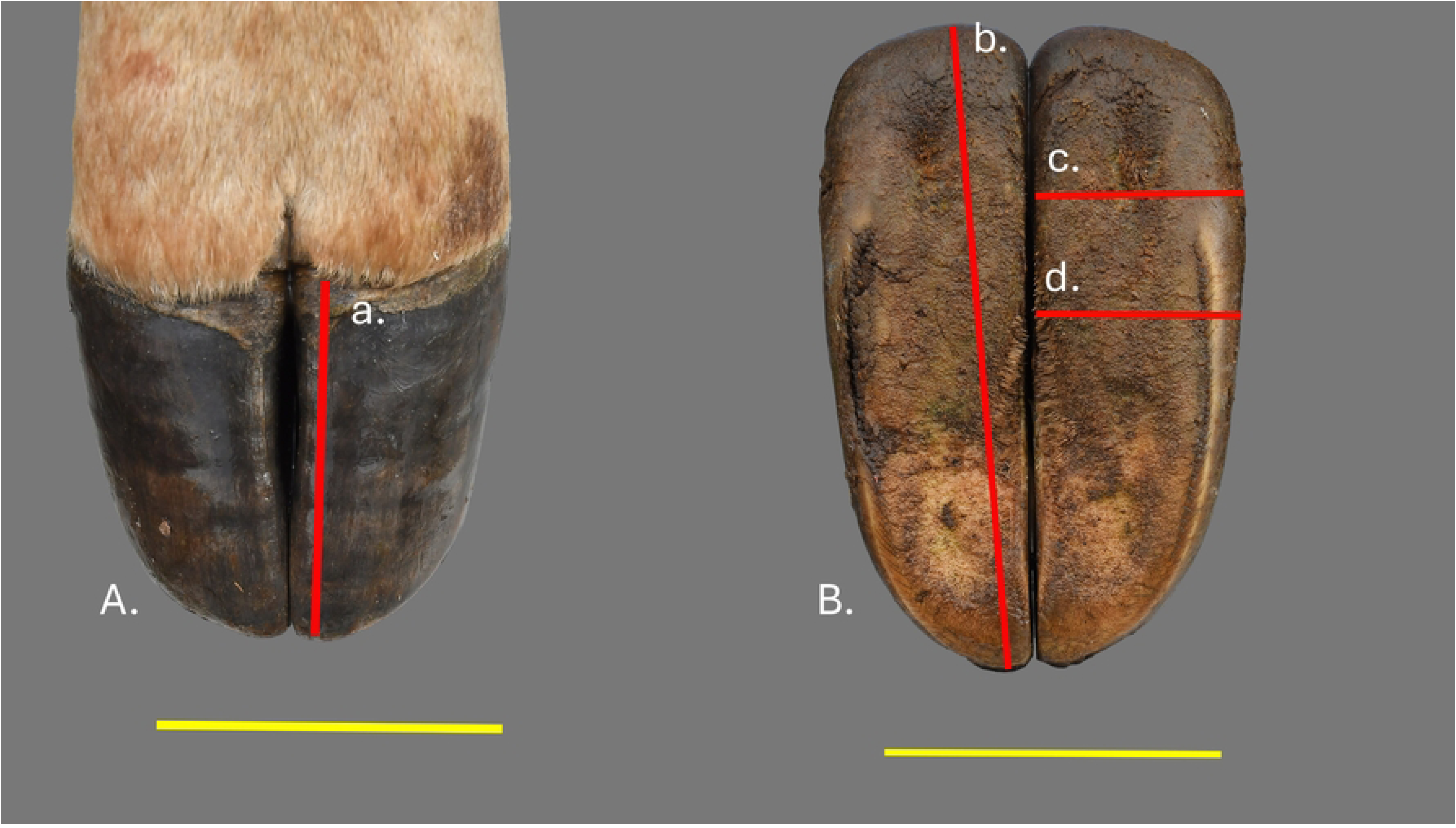
Right thoracic foot of a wild Angolan giraffe. (A) Dorsal view. (B) Distal view. a-toe height. b-claw length. c- heel width. d- median width. Scale bar equals 10cm.

**Figure 6.**
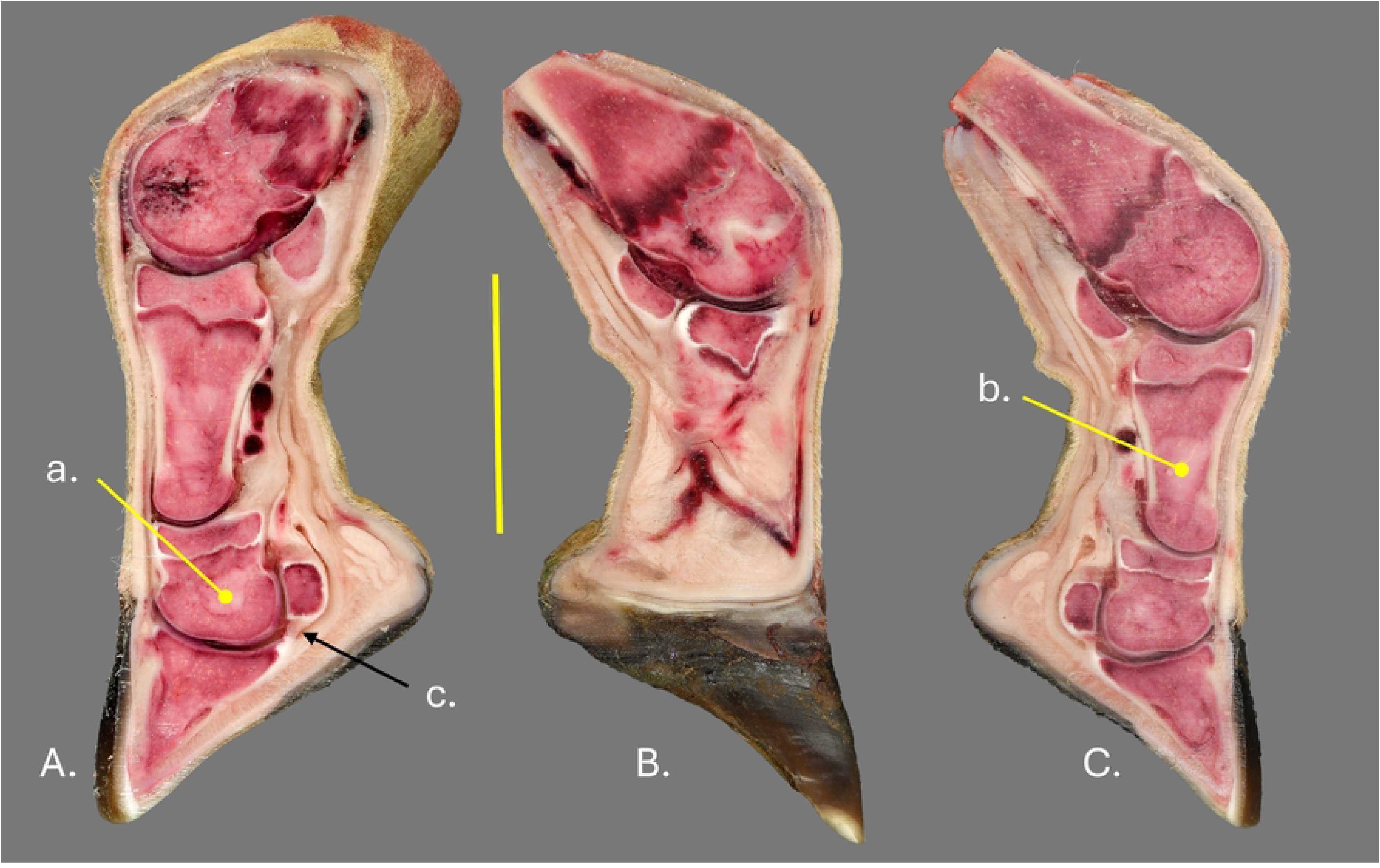
Left thoracic distal limb of a giraffe in human care. (A) Sagittal section through lateral claw. (B) Mid-sagittal section. (C) Sagittal section through the medial claw. a- bone marrow in middle phalanx. b- bone marrow in proximal phalanx. Scale bar equals 10cm.

Anatomical dissections were made using right limbs. Prior to dissection, the hoof capsules were removed using a bone saw, chisel and blunted hoof nippers. Vertical cuts were made on the abaxial hoof wall at the toe and heel and a cut was made on the distal surface of the foot just inside the white line. The abaxial wall was then pried away using the chisel and hoof nippers.

Lastly, the sole, axial wall and heels were also removed using the chisel and hoof nippers. After removal of the hoof capsule, some limbs were dissected to demonstrate the vasculature while others were dissected to demonstrate tendons and ligaments. Dissection proceeded through the disarticulation of the metacarpophalangeal and interdigital joints. Photos and notes were taken frequently during dissection to record anatomical findings. It was not possible to preserve the bones from the wild individuals, and they were disposed of as per ethics recommendations.

Selected human care specimens were macerated using dermestid beetles and soaked in 7% hydrogen peroxide. These disarticulated bones were then used to inform the detailed descriptions below.

## Results

### Bones

The description presented includes anatomy from the distal metapodial down. The metapodials (Fig 7) consist of fused 3^rd^ and 4^th^ metatarsals/metacarpals and are very robust elements with thick cortical bone and a well-defined medullary cavity. The metacarpal is proportionally more robust than the metatarsal. The head of the metapodial is divided into two trochlea by a deep intertrochlear notch (Fig 7a). Each trochlea is further divided by a median ridge extending across the entire articular surface of the trochlea (Fig 7b). Deep fossae on the abaxial surface of the trochlea provide attachment for strong collateral ligaments. Two proximal sesamoids articulate with the palmar/plantar surface of each trochlea on either side of the median ridge. The axial proximal sesamoids are flatter than the abaxial ones (Fig 8).

**Figure 7.**
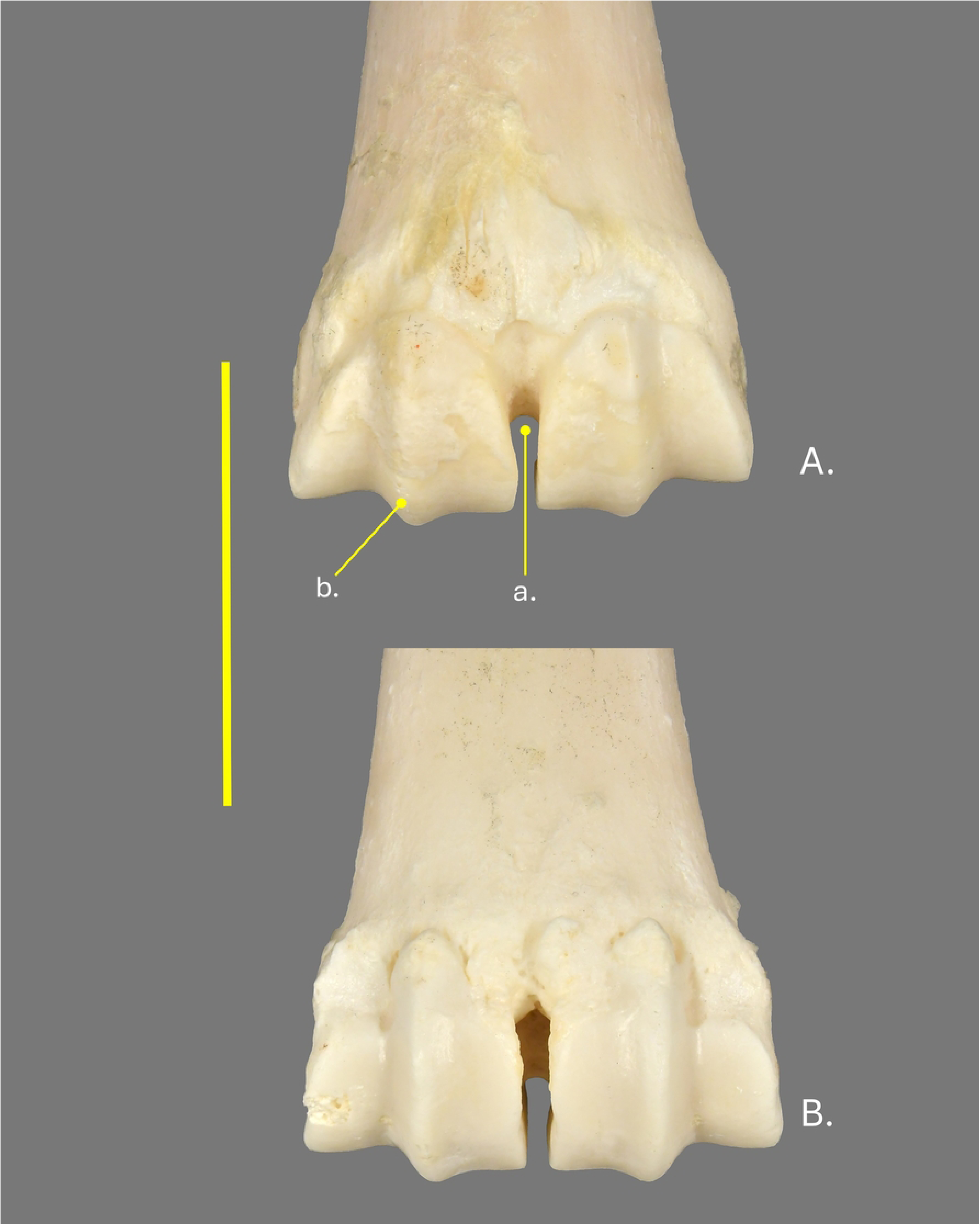
Right distal metacarpal of a giraffe in human care. (A) Dorsal view. (B) Palmar view. a- inner trochlear notch. b- median intratrochlear notch. Scale bar equals 10cm.

**Figure 8.**
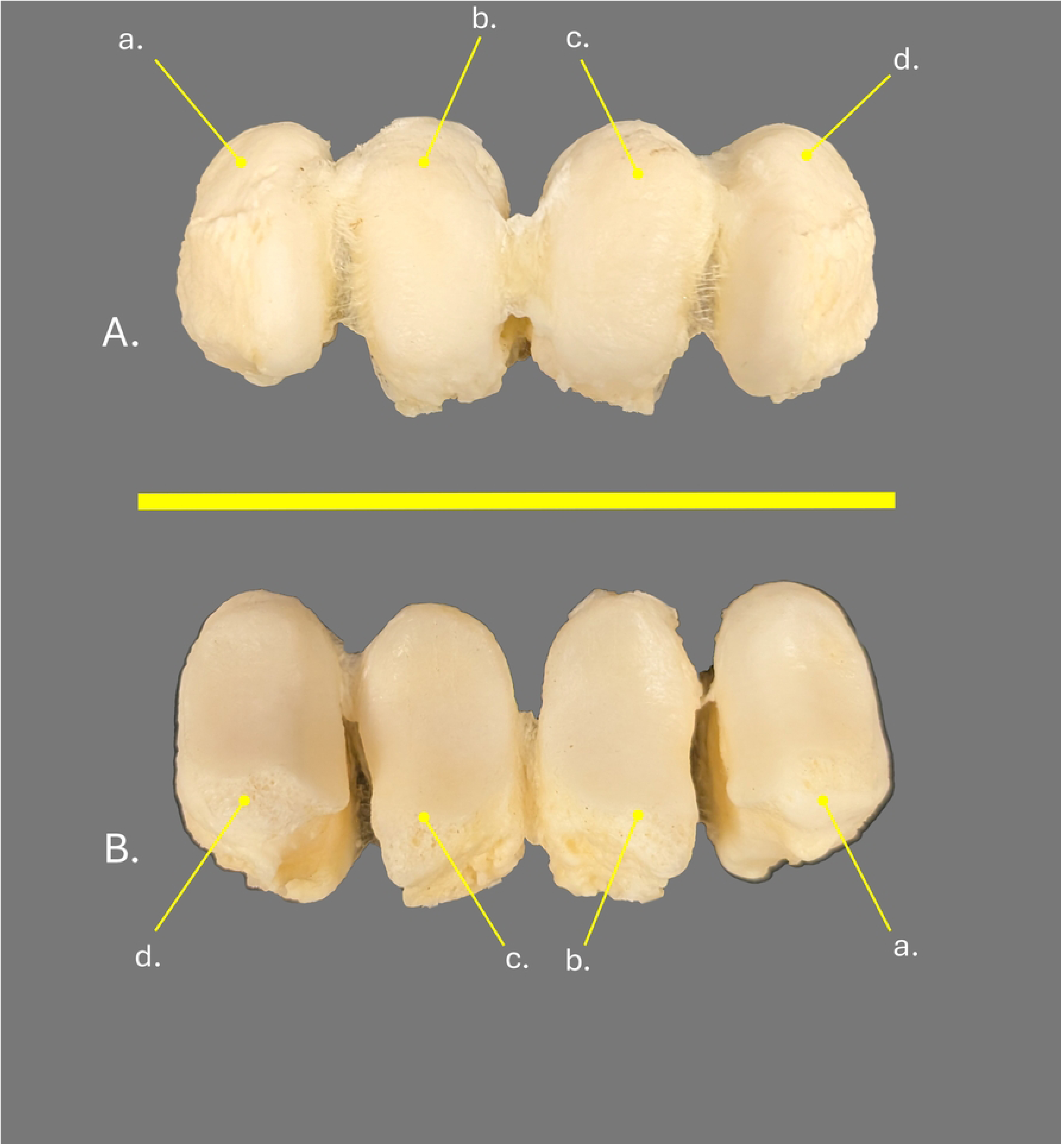
Right thoracic limb proximal sesamoids of a giraffe in human care. (A) Palmar view. (B) Dorsal view. a- medial abaxial proximal sesamoid. b- medial axial proximal sesamoid. c- lateral axial proximal sesamoid. d- lateral abaxial proximal sesamoid. Scale bar equals 10cm.

Digits three and four are present in giraffe with both consisting of three phalanges. All three phalanges have discernible medullary cavities (Fig 1) with those of the 2.5 month containing more red marrow (Fig 6) than the older specimens which all appear to have yellow marrow. The medullary cavity in the proximal and middle phalanges are very prominent and filled with adipose tissue in all the specimens examined while the medullary cavity the distal phalanx is visible but has less defined endosteal edges than the medullary cavities of both the middle and distal phalanges (Fig 1).

The proximal phalanx (P1) (Fig 9) is the longest of the three and has a flattened axial margin and curved abaxial margin. Two prominent ligament scars for attachment of the distal sesamoidean ligaments are present on the proximal palmar/plantar surface of the proximal phalanx. The axial ligament scar (Fig 9a) is significantly longer and more robust than the abaxial ligament scar (Fig 9b). The trochlea on the distal articular surface of the proximal phalanx is divided into two parts by a deep groove with the abaxial area of the articular surface being significantly larger than that of the axial portion (Fig 9c). Deep fossae are present on the distal end of the proximal phalanx for the attachment of the collateral ligaments of the proximal interphalangeal joint (Fig 9d).

**Figure 9.**
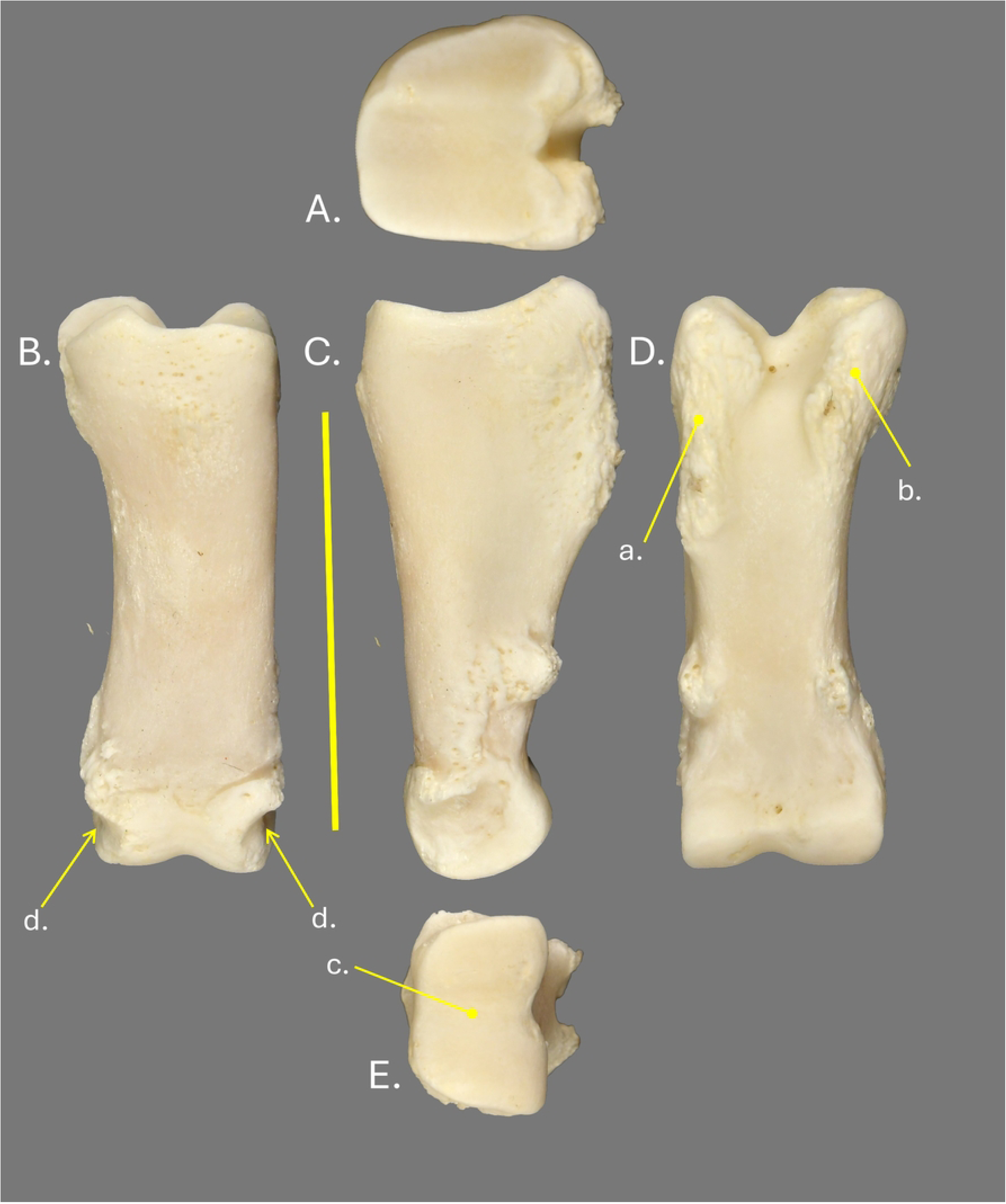
Right lateral proximal phalanx of a giraffe in human care. (A) Proximal view. (B) Dorsal view. (C) Medial view. (D) Palmar view. (E) Distal view. a- axial sesamoidian ligament scar. b- abaxial sesamoidian ligament scar. c- distal intratrochlear groove. d- collateral ligament attachments. Scale bar equals 10cm.

The middle phalanx (P2) (Fig 10) is short and robust. The proximal articular surface mirrors the proportions of the trochlea of the proximal phalanx with a ridge dividing the articular surface (Fig 10a). A prominent muscle scar is present on the proximal dorsal surface of the middle phalanx for the attachment of extensor tendons (Fig 10b). The distal end of the middle phalanx is significantly asymmetric with the articular trochlea divided by a deep groove that runs at an oblique angle from the dorsal abaxial surface to the center of the palomar/plantar surface (Fig 10c). The groove separates a small abaxial articular surface from a much larger axial articular surface. This asymmetry results in the axial separation of the distal phalanges. A shallow fossa is present on the abaxial surface of the middle phalanx for the attachment of the collateral ligament of the distal interphalangeal joint.

**Figure 10.**
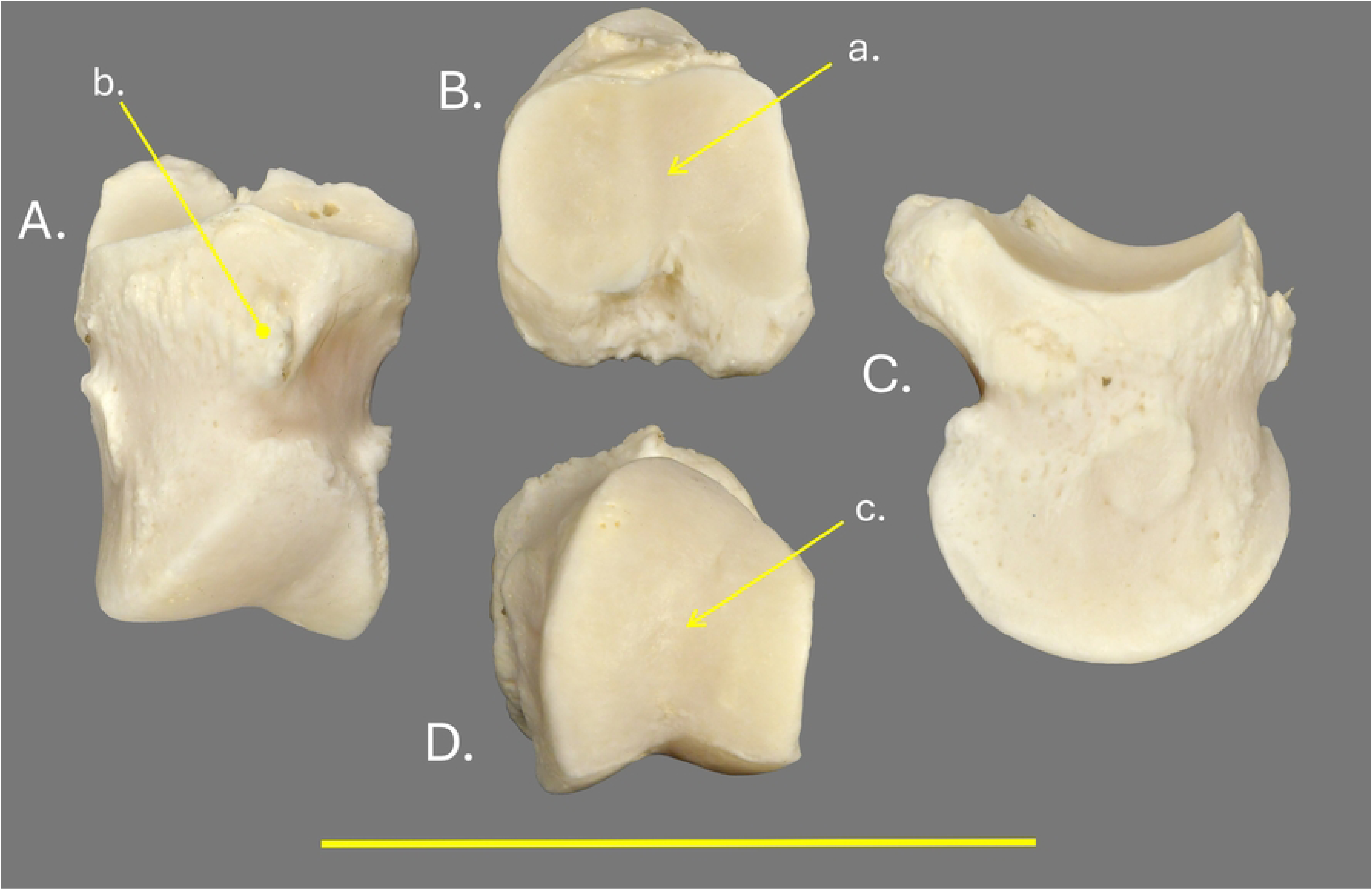
Right lateral middle phalanx of a giraffe in human care. (A) Dorsal view. (B) Proximal view. (C) Medial view. (D) Distal view. a- dorsal median ridge b- digital extensor attachment scar. c- intratrochlear groove. Scale bar equals 10cm.

The distal phalanx (P3) (Fig 11) is extremely asymmetric with the axial side being virtually straight while the abaxial surface presents an arched surface extending from the tip of the distal phalanx to the level of the dorsal margin of articular surface with the middle phalanx. The articular surface of the distal phalanx mirrors that of the trochlea of the middle phalanx with a ridge dividing the much larger abaxial portion from the small axial portion of the articular surface (Fig 11a). The entire articular surface is also angled such that the abaxial articular surface is farther from the distal surface of the bone than the axial surface. A prominent flattened area on the palmar/plantar region just distal to the articular surface defines the broad flexor tubercle for the insertion of the deep digital flexor tendon (Fig 11b). The extensor process is formed by a broad, roughly triangular area on the dorsal apex of the distal phalanx (Fig 11c). Large foramina on both the abaxial (Fig 11d) and axial surface (Fig 11e) of the distal phalanx just below the articular surface for passage of the arcuate vessels. The distal surface of the distal phalanx is perforated by several large solar foramina for vessels supplying the solar dermis (Fig 11f). Small, parallel bony axial to abaxial ridges may be seen in older individuals, corresponding to the septa between compartments of the digital cushion.

**Figure 11.**
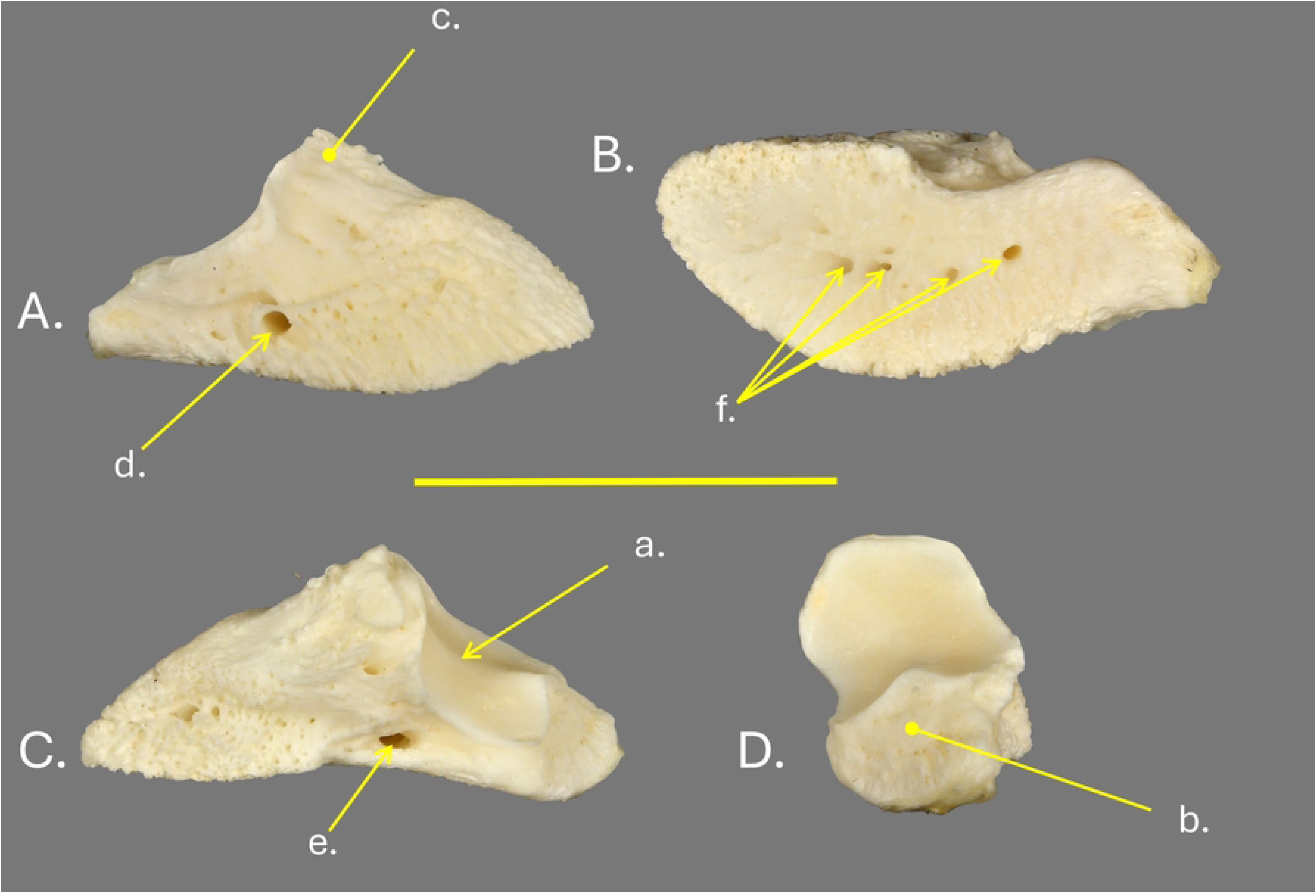
Right lateral distal phalanx of a giraffe in human care. (A) Lateral view. (B) Distal view. (C) Medial view. (D) Proximal view. a- dorsal median ridge. b- flexor tubercle. c-extensor process. d- abaxial foramen for arcuate artery. e- axial foramen arcuate artery. f- solar foraminia. Scale bar equals 10cm.

The distal sesamoids (Fig. 12) are roughly rectangular with an expanded palmar/plantar surface which corresponds to the width of the distal deep digital flexor tendon. The dorsal surface is formed by the articulates with the palmar/plantar of the middle phalanx (Fig 1e). The articular surface is divided by a ridge (Fig 12a) into a large lateral articular surface and a smaller medial articular surface.

**Figure 12.**
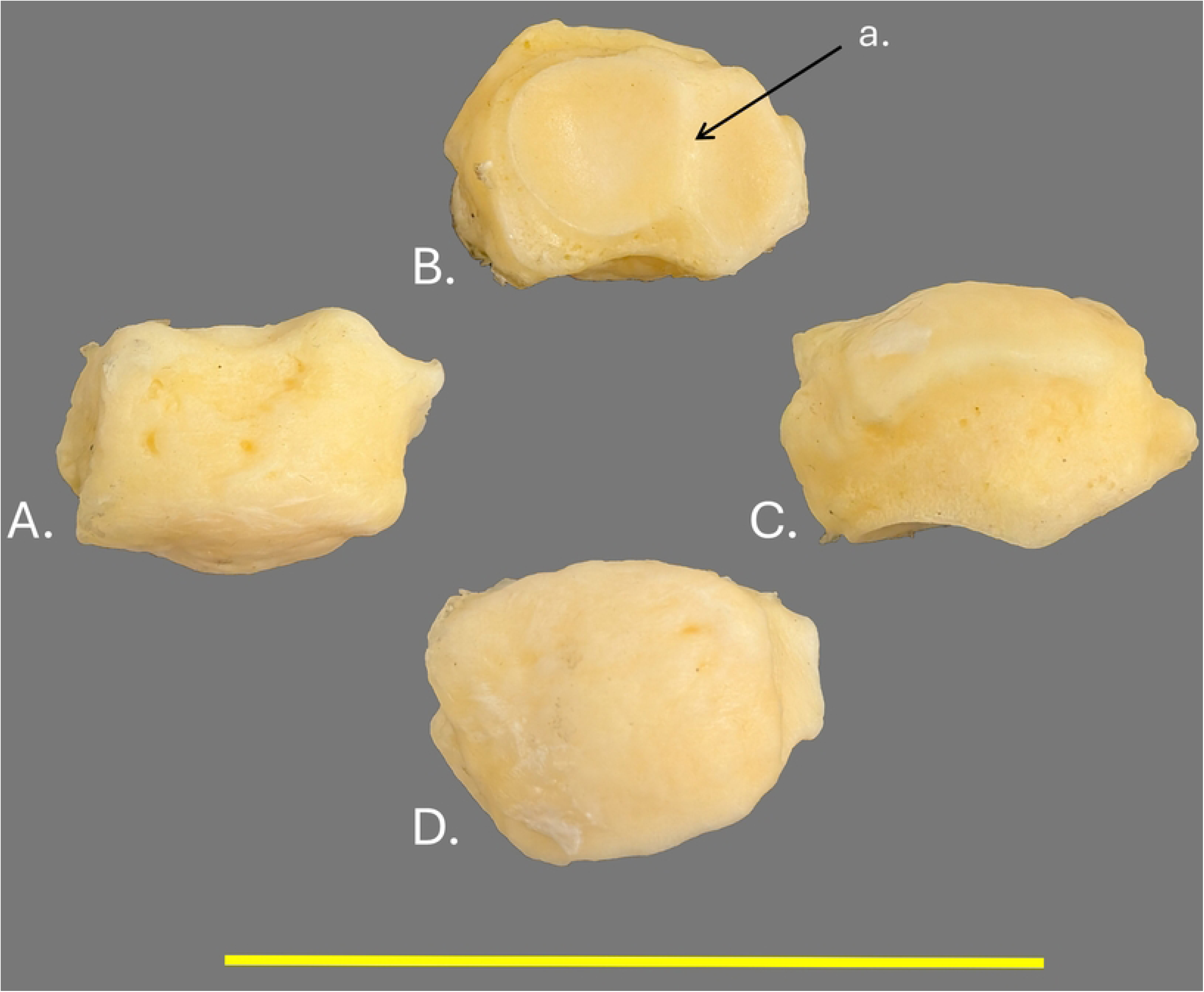
Right medial distal sesamoid of a giraffe in human care. (A) Proximal view, (B) Dorsal view, (C) Distal View (D) Palmar view. Scale Bar equals 10cm.

### Ligaments of the Joints

A thick, strong proximal interdigital ligament is found between the axial palmar/plantar proximal surfaces of the proximal phalanges (Fig 13a). Collateral ligaments are found at each joint and will be described from proximal to distal. The metacarpophalangeal/metatarsophalangeal joint has two sets of collateral ligaments corresponding to the articulations of digits three and four. The abaxial collateral ligaments of the metacarpophalangeal/metatarsophalangeal joints are by far the most robust with a large attachment to the deep fossae on the abaxial sides of the metapodial and a broad attachment to the abaxial proximal surface of the proximal phalanx (Fig 14a). The axial collateral ligaments of the metacarpophalangeal/metatarsophalangeal joints are much smaller and the attachment to the metapodial is deep in the intertrochlear notch. The narrow ligament then widens into a thin broad fan-shaped attachment to the proximal palmar/plantar axial surface of the proximal phalanx (Fig 14b).

**Figure 13.**
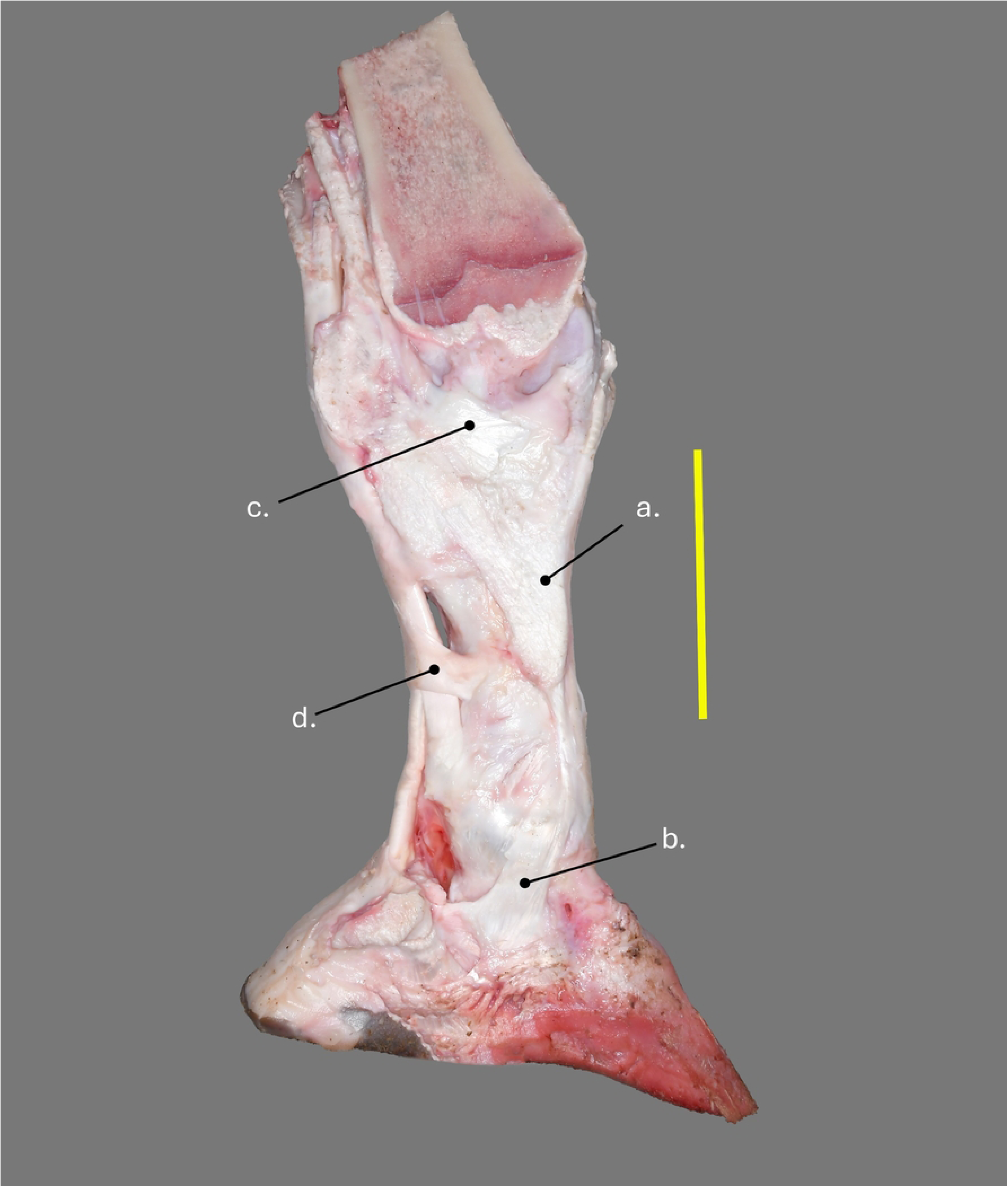
Right thoracic medial claw in axial view of a wild Angolan giraffe. a- proximal interdigital ligament. b- axial collateral ligament of the distal interphalangeal joint. c- axial extensor branch of the interosseous. d- proximal digital annular ligament. Scale bar equals 10cm.

**Figure 14.**
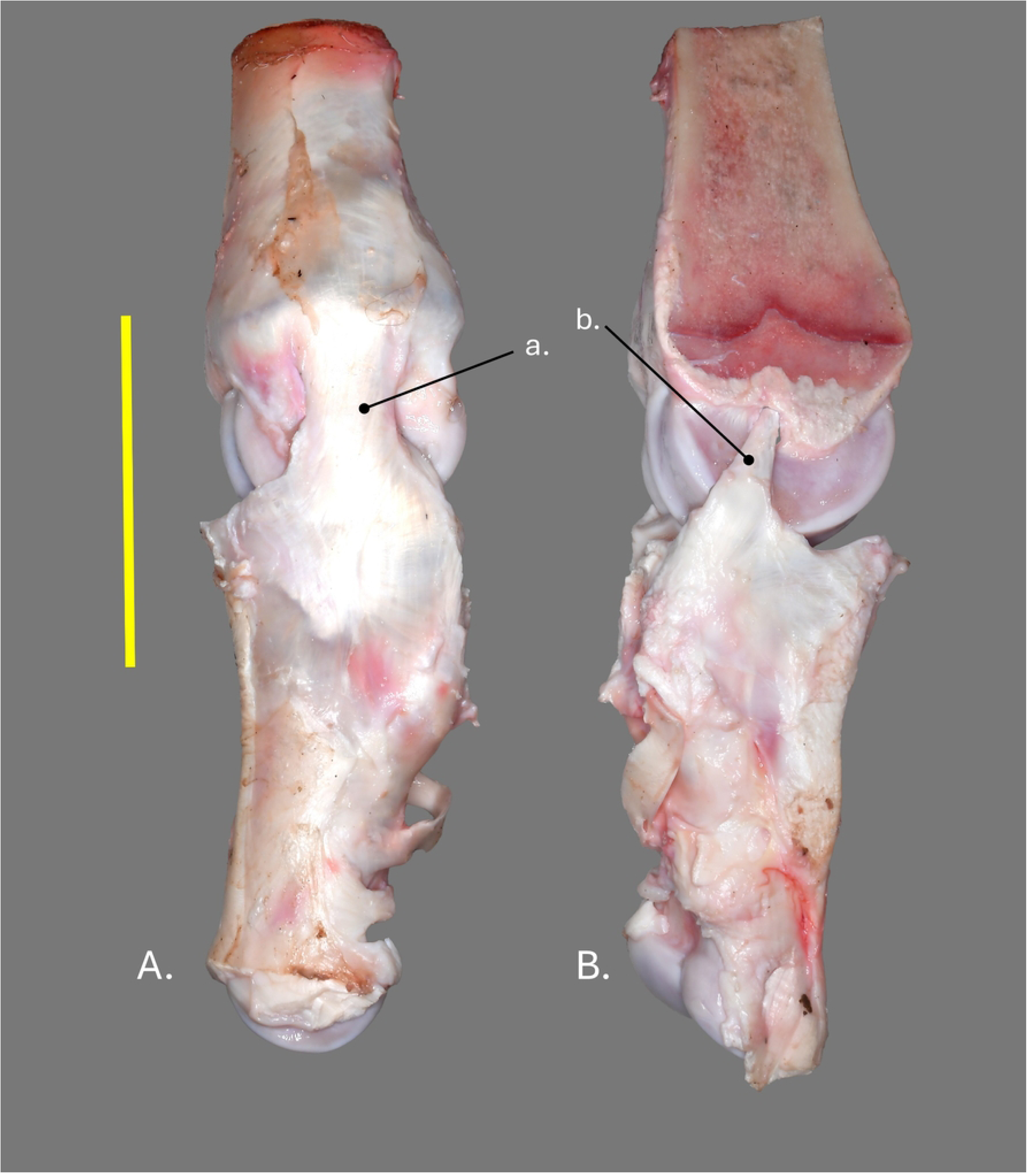
Right thoracic medial digit of a wild Angolan giraffe. (A) Abaxial view. (B) Axial view. a- abaxial collateral ligament of the metacarpophalangeal joint. b- axial collateral ligament of the metacarpophalangeal joint. Scale bar equals 10cm.

The collateral ligaments of the proximal interphalangeal joint are more complex. On the axial side, a superficial part of the ligament is attached proximally to the distal axial border of the proximal phalanx. This superficial portion of the ligament then forms a thin broad ligament extending distally to attach to the distal sesamoid, axial border of the distal phalanx and finally to the dense connective tissue encapsulating the digital cushion (Fig 13b). A similar pattern is repeated on the abaxial side. The deep components of the collateral ligaments of the proximal interphalangeal joint consists of stout axial and abaxial ligaments which attach to deep fossae on either side of the distal proximal phalanx. Each ligament then runs caudodistally to attach to the proximal palmar/plantar surface of the middle phalanx mixing with the tendon of insertion of the superficial digital flexor (Figure 15 a and b).

**Figure 15.**
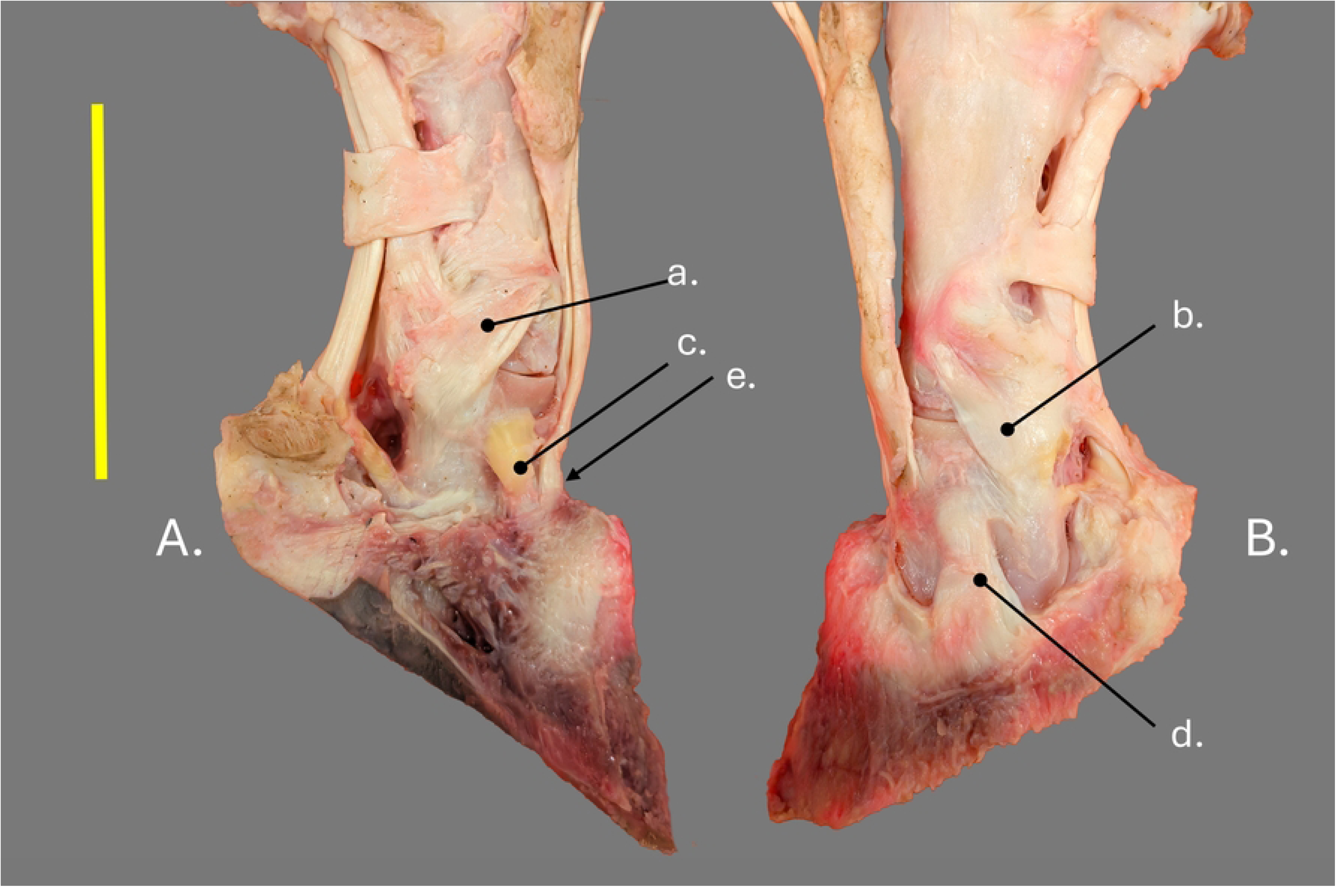
Right pelvic limb medial claw of a giraffe in human care. (A) Axial view. (B) Abaxial view. a- axial collateral ligament of the proximal interphalangeal joint. b- abaxial collateral ligament of the proximal interphalangeal joint. c- axial collateral ligament of the distal interphalangeal joint. d- abaxial collateral ligament of the distal interphalangeal joint. e- long digital extensor tendon. Scale bar equals 10cm.

The abaxial collateral ligament of the distal interphalangeal joint is very stout and attaches proximally in a deep fossa on the distal abaxial side of the middle phalanx (Fig 15d). The ligament then broadens somewhat distally where it is attached to the abaxial proximal border of the distal phalanx directly adjacent to the joint capsule. The axial collateral ligament is very robust and consists primarily of elastic connective tissue as indicated by its yellow color (Fig 15c). The axial collateral ligament is attached proximally to a prominent ligament scar found on the axial side of the middle phalanx. The ligament then runs for a short distance to attach to the axial side of the extensor process of the distal phalanx where it merges with the attachment of the common/long digital extensor tendon (Fig 14e).

A very short stout impar ligament connects the distal phalanx to the distal sesamoid on each digit. It is most easily seen in cross sectional view (Fig 6c).

### Tendons

The tendons of the distal limb are described beginning at the mid metapodial. The tendons of the common/long digital extensor and lateral digital extensor merge to form a stout narrow tendon extending down the dorsal surface of the thoracic limb. In the pelvic limb, the long digital extensor and lateral digital extensor merge in a similar way. In both the thoracic and the pelvic limbs the combined tendon of the digital extensors separate into distinct branches just above the fetlock joint (Fig 16). Two broad abaxial branches lie on the dorsal aspect of the proximal phalanx where they cross the proximal interphalangeal joints and insert on the dorsal proximal surface of the middle phalanges (Fig. 16a and b). Lying between these abaxial branches, a single stout round common tendon courses over the dorsal surface of the fetlock joint (Fig 16c). As this common tendon passes over the fetlock joint, it splits into two proper axial branches which run in a digital extensor tendon sheath from the level of the fetlock to the insertion of the tendons onto the extensor process of the distal phalanx (Fig 16 d and e). These two tendons are much smaller than the abaxial branches and are oval in cross section (Fig 17b).

**Figure 16.**
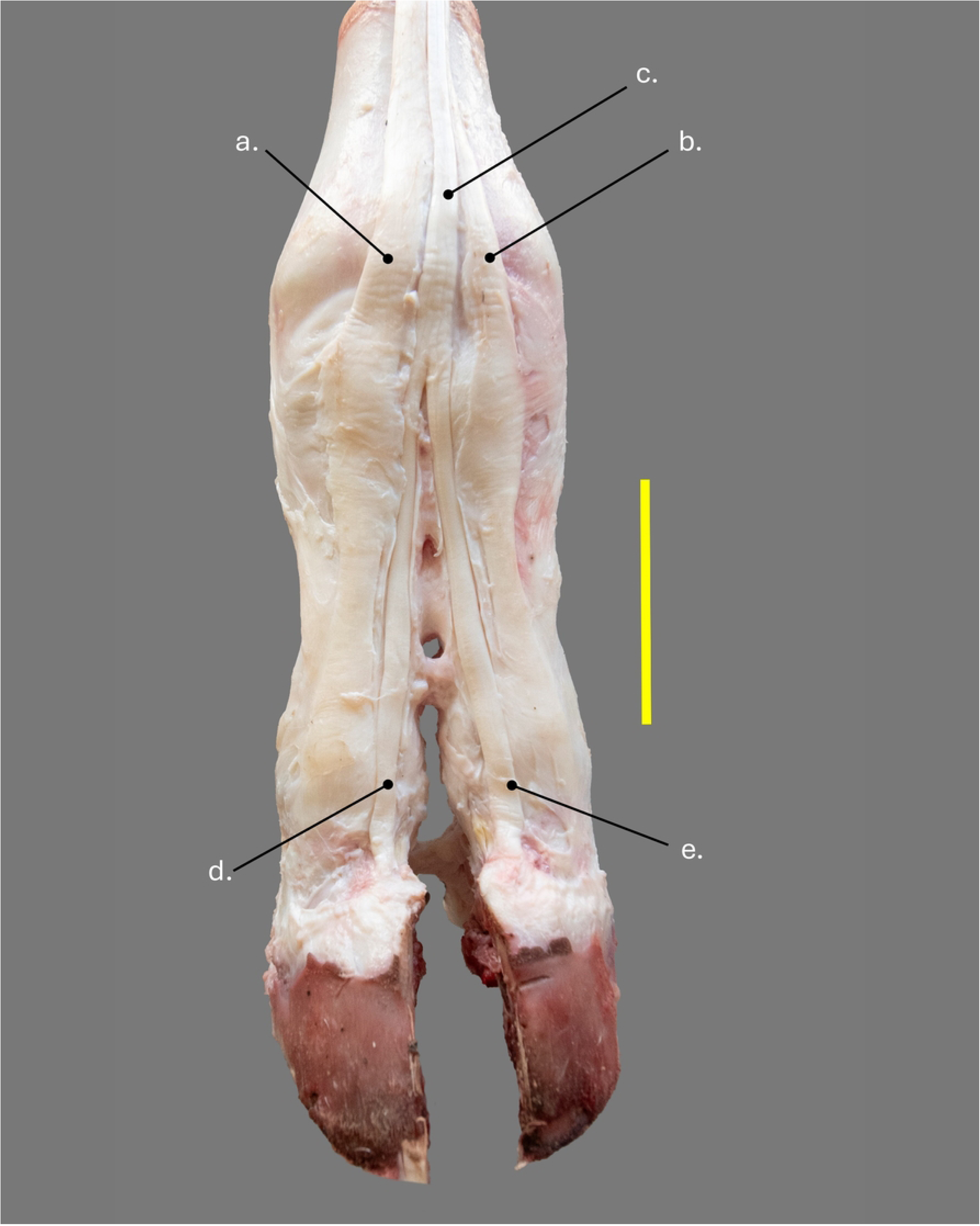
Right pelvic limb in dorsal view of a wild Angolan giraffe. a- lateral extensor tendon. b- medial extensor tendon. c- middle extensor tendon. d- axial proper digital extensor tendon for digit IV. e- axial proper digital extensor tendon for digit III. Scale bar equals 10cm.

**Figure 17.**
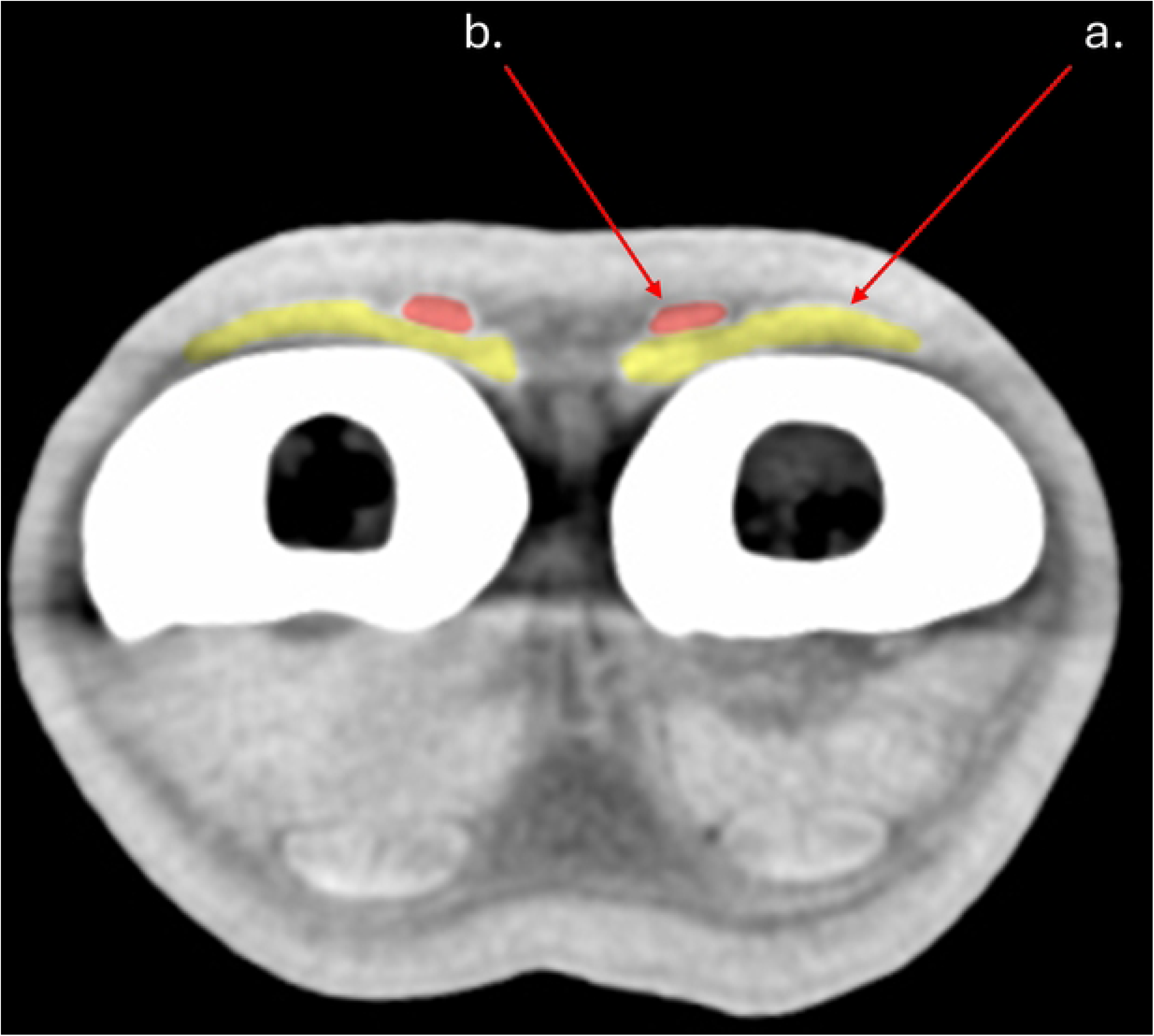
CT mid-sagittal section through the left pelvic limb of a wild Angolan giraffe. a-middle extensor tendon. b- axial proper digital extensor tendon.

The superficial digital flexor, deep digital flexor and interosseus tendons have virtually identical morphologies in both the thoracic and pelvic limb and the description that follows applies to both. The deep digital flexor is a thick wide oval tendon in the middle of the metapodial lying between the interosseus and the superficial digital flexor (Figs 3c and 18b). Just above the fetlock joint, the deep digital flexor divides into distinct medial and lateral branches. Both branches widen as they cross the level of the proximal sesamoids. Distal to the proximal sesamoids the tendon narrows continuing distally. At the level of the distal sesamoids, each tendon widens dramatically covering the entire distal surface of the distal sesamoid and separated from the sesamoid by a navicular bursa. Each tendon then inserts on the flexor tubercle which is formed by the entire palmar/plantar surface of the distal phalanx. Each branch of the deep digital flexor tendon lies in a digital flexor tendon sheath beginning at the level of the fetlock joint and ending just as the tendon begins to widen over the distal sesamoid. (Fig 18).

**Figure 18.**
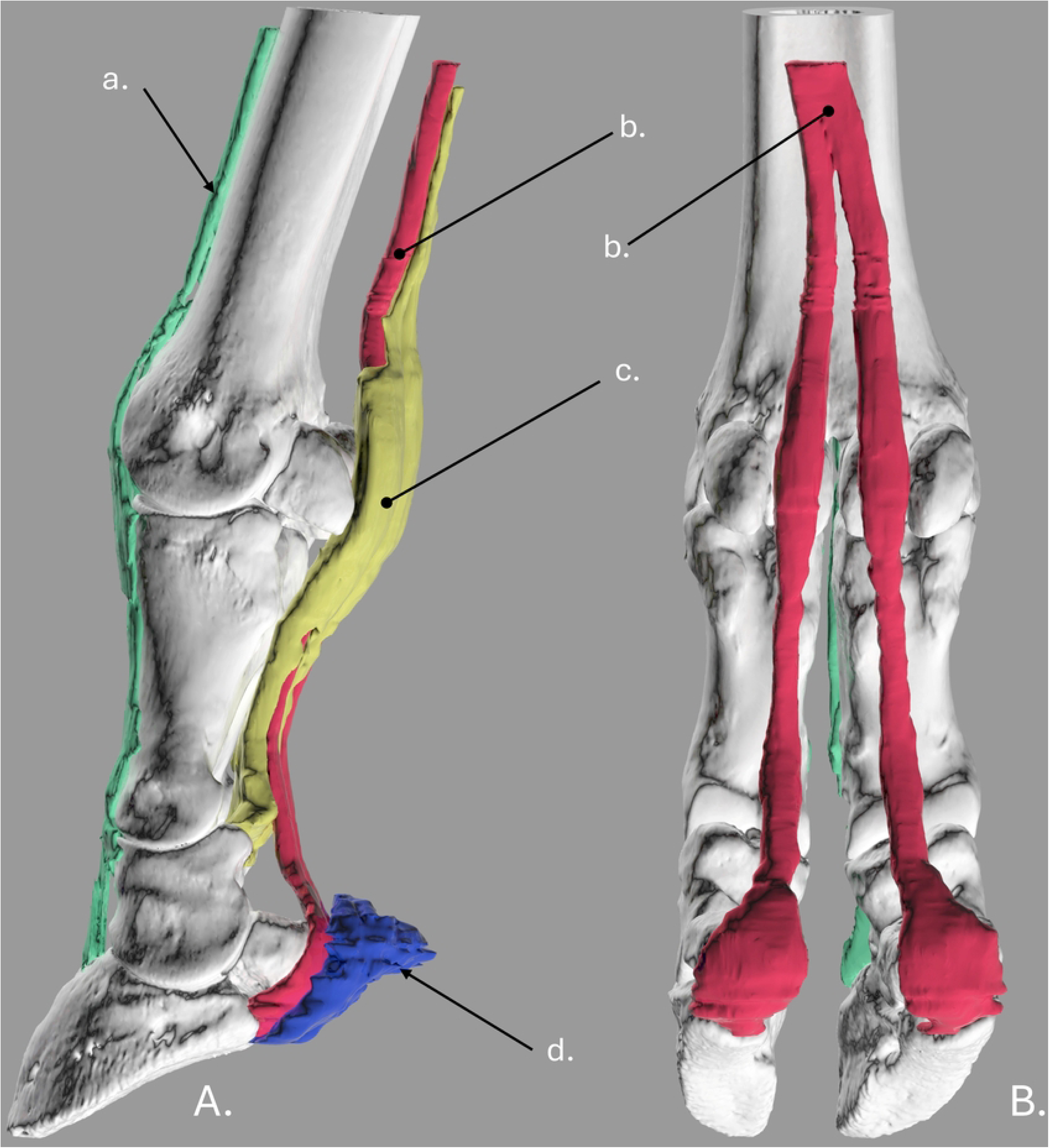
3-D reconstruction of the right distal thoracic limb of a wild Angolan giraffe. (A) Medial view. (B) Palmar view. a- digital extensors. b- deep digital flexor. c- superficial digital flexor. d- adipose tissue of the heel bulb.

The superficial digital flexor covers the deep digital flexor above the fetlock. Just proximal to the fetlock, the superficial digital flexor splits into two branches. Each branch widens before it crosses the proximal sesamoids and totally surrounds the deep digital flexor to form the flexor manica (Fig 19c). The flexor manica is open on its deep surface such that the deep digital flexor is visible at the level of the proximal sesamoids (Fig 20a). Distal to the proximal sesamoids, the superficial digital flexor tendons lie deep to the deep digital flexor tendons and are wide and thick. Each tendon then inserts onto the proximal palmar/plantar surface on the middle phalanx just behind the joint capsule (Fig 19c).

**Figure 19.**
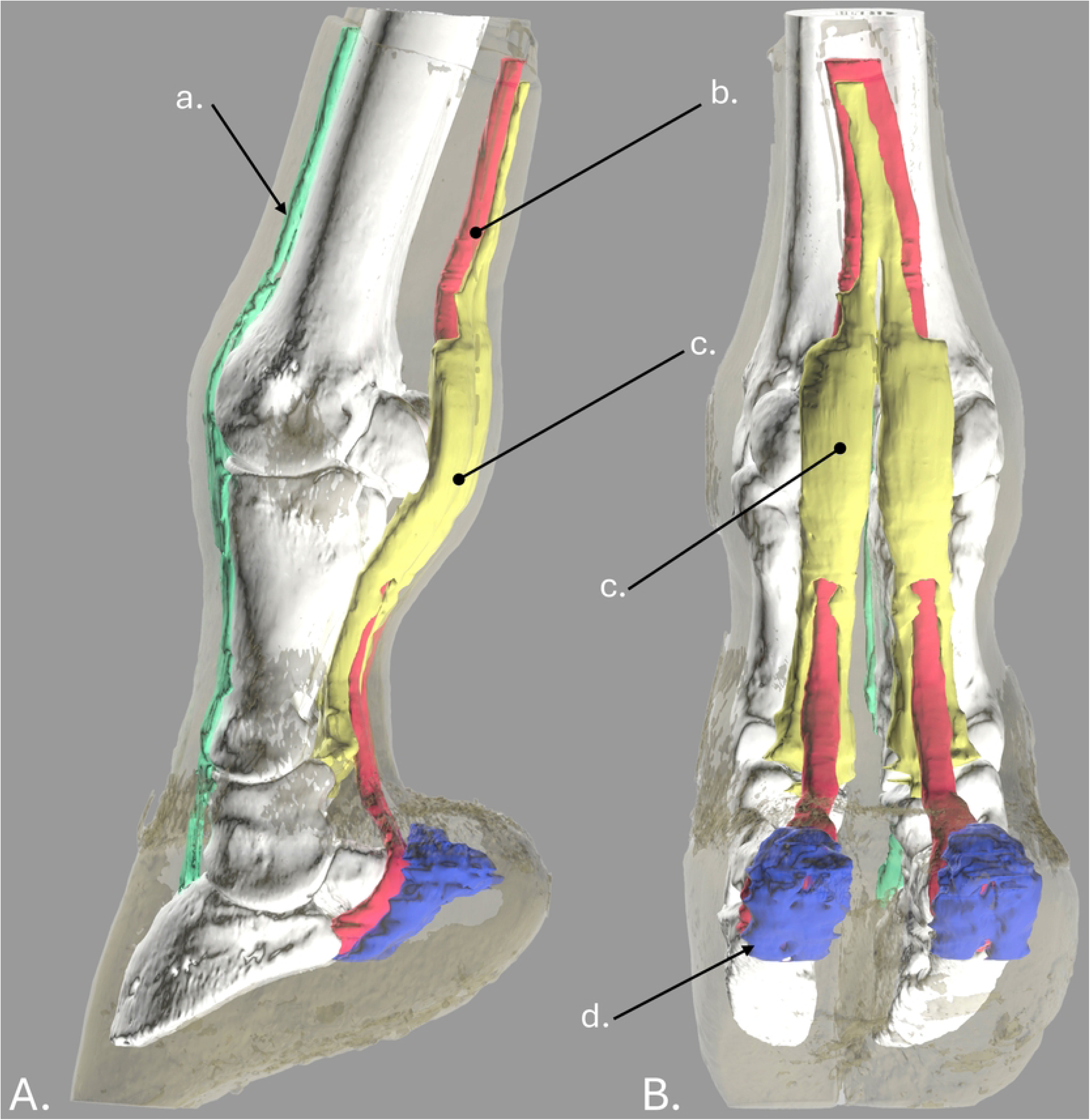
3-D reconstruction of the right distal thoracic limb of a wild Angolan giraffe. (A) Medial view. (B) Palmar view. a- digital extensors. b- deep digital flexor. c- superficial digital flexor. d- adipose tissue of the heel bulb.

**Figure 20.**
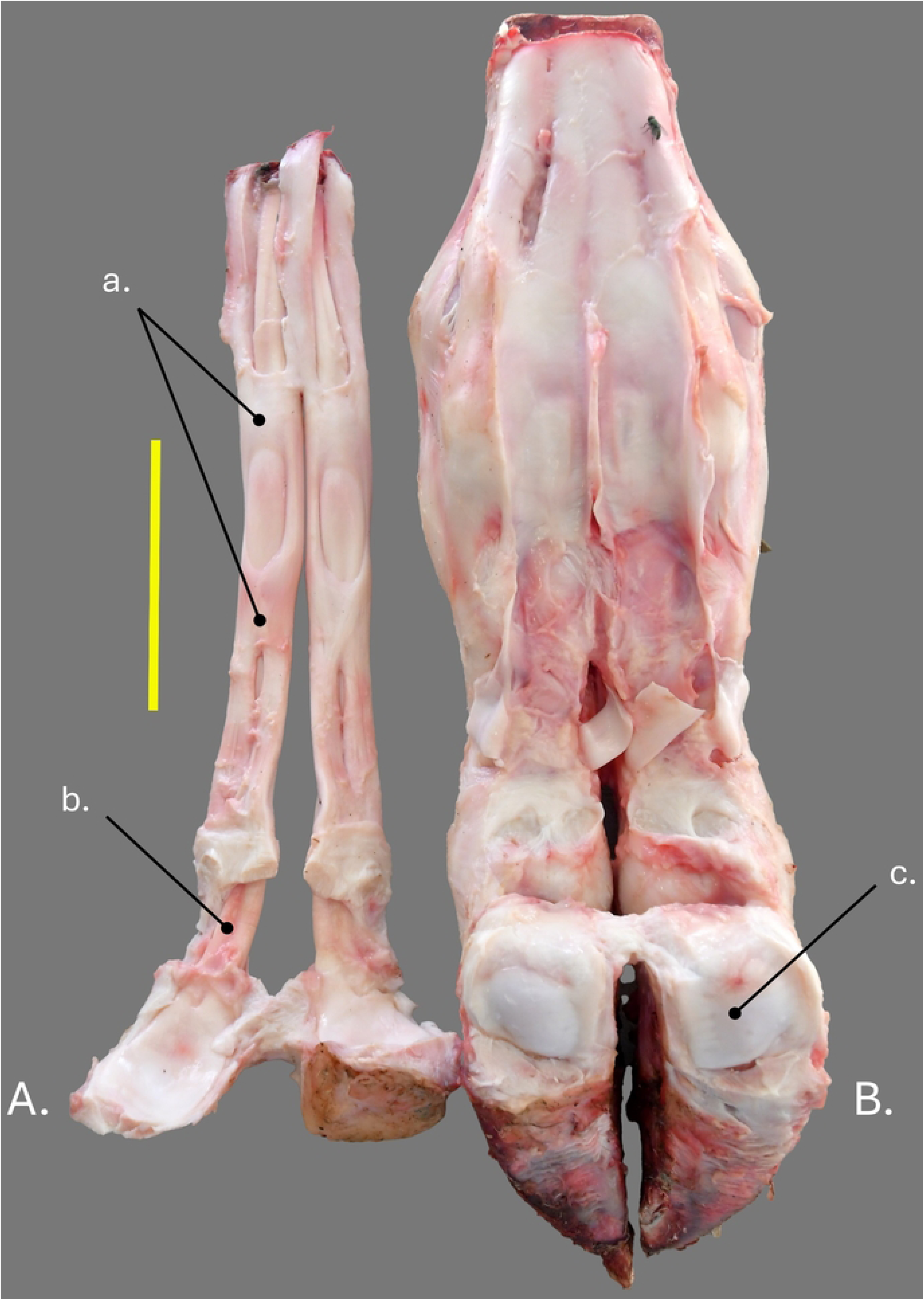
Right thoracic distal limb of a wild Angolan giraffe. (A) Combined superficial and deep digital flexors in dorsal view. (B) Palmar view with superficial and deep digital flexors removed. a- superficial digital flexor. b- deep digital flexor. c- distal sesamoid (navicular bone). Scale bar equals 10cm.

The interosseus in the distal metapodial region consists of a broad thick tendon which lies against the palmar/plantar surface of the metapodial. About 10 cm above the fetlock, the tendon divides into four primary branches that insert onto each of the proximal sesamoids (Fig 21a).

**Figure 21.**
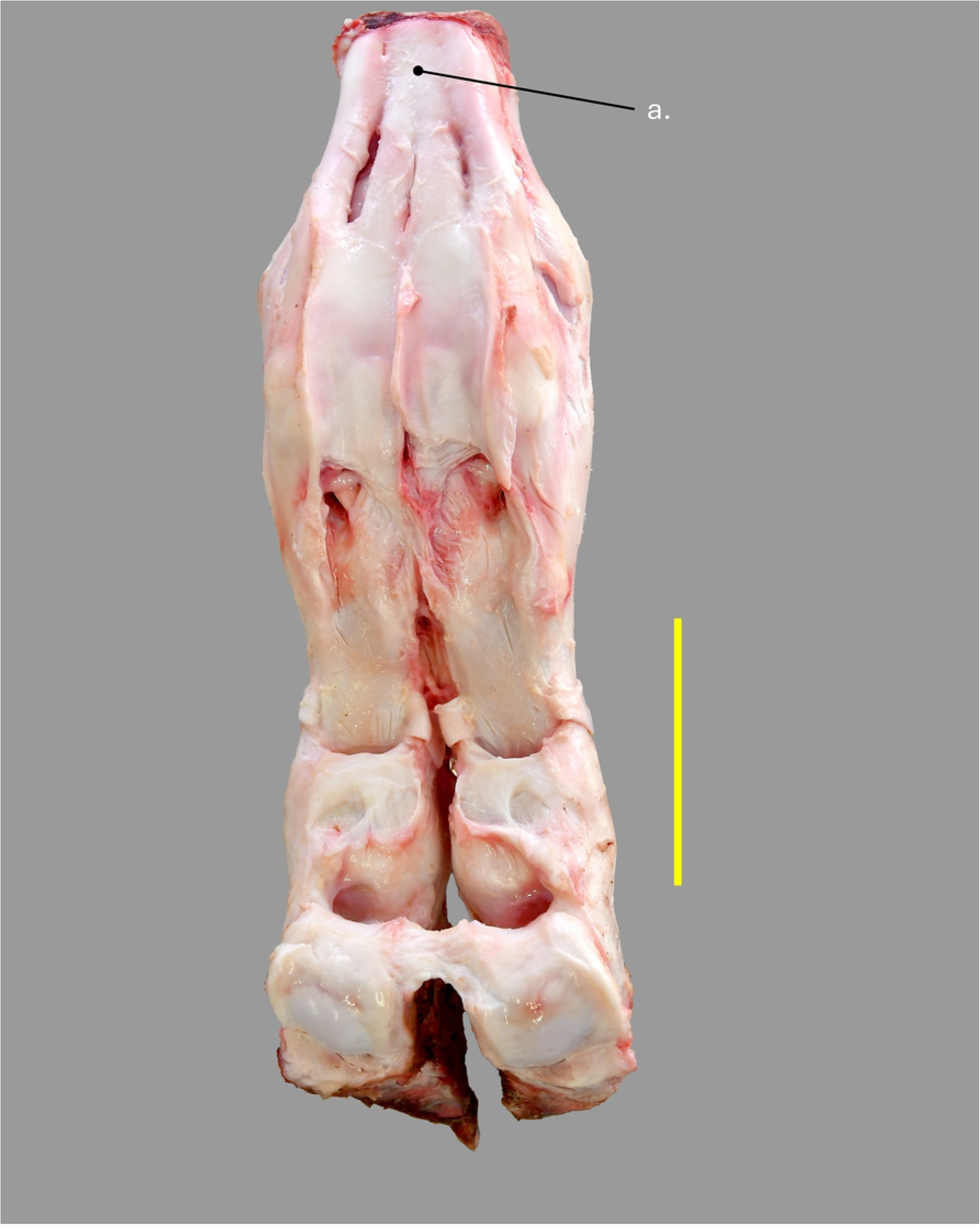
Right thoracic distal limb in palmar view with superficial and deep digital flexors removed of a wild Angolan giraffe. a- interosseous (suspensory ligament). Scale bar equals 10cm.

Abaxial extensor branches extend from the interosseus and wrap around the dorsal surface of the proximal phalanx to unite with the broad abaxial digital extensor branches of digits III and IV. A common stout axial extensor branch courses between the axial proximal sesamoids and helps form the proximal interdigital ligament (Fig 13c). This single axial branch then splits into two axial extensor branches which wrap around the axial surface of each digit to unite with the abaxial digital extensor branches of digits III and IV.

### Annular Ligaments

A thin palmar annular ligament binds down the tendons of the superficial and deep digital flexors. The palmar annular ligament is attached to the palmar/plantar surface of both the axial and abaxial proximal sesamoids (Fig 22a). A broad stout proximal digital annular ligament binds down the superficial and deep digital flexor tendon and attaches to the axial and abaxial distal surface of each proximal phalanx (Figs 13d and 22b). The distal digital annular ligaments are complex and combine to form the distal interdigital ligament (Fig 22c and d). Each distal digital annular ligament runs at an angle from proximal on the abaxial side to distal on the axial side of digits III and IV. The ligament has its abaxial attachment to both the distal abaxial palmar/plantar surface of the proximal phalanx and the proximal abaxial palmar/plantar surface of the middle phalanx. The axial portion of the distal digital annular ligament is attached to the axial border of the distal sesamoid. Transverse fibers also connect the axial surfaces of the distal digital annular ligament to form the distal interdigital ligament. The distal interdigital ligament is very strong and is located between the proximal borders of the heel bulbs (Fig 22d).

**Figure 22.**
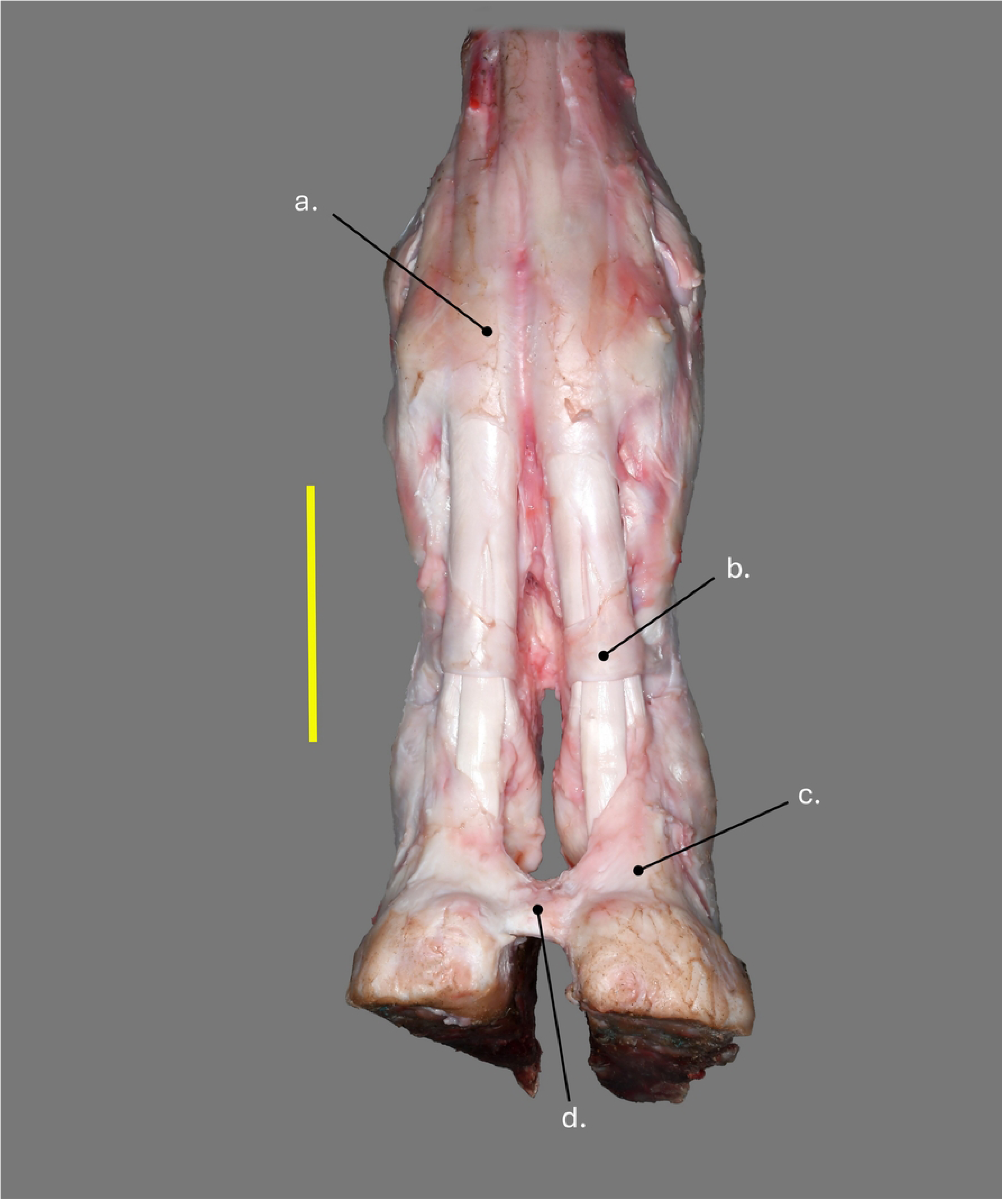
Right thoracic distal limb in palmar view of a wild Angolan giraffe. a- palmar annular ligament. b- proximal digital annular ligament. c- distal digital annular ligament. d- distal interdigital ligament. Scale bar equals 10cm.

### Arteries of the Front Foot

At the level of the fetlock, a large palmar common digital artery III runs on the axial palmar surface of the foot and is the largest artery supplying the foot. It begins superficial to and in between the palmar annular ligaments and then courses down the axial interdigital space where it moves deeper into the interdigital space (Fig. 23a). Near the level of the distal end of the proximal phalanx, the artery divides into axial palmar proper digital artery III supplying the third digit and axial palmar proper digital artery IV supplying the fourth digit (Fig 23b and c). Each axial palmar proper digital artery then gives an abaxial palmar proper digital artery to its respective digit. One branch of each abaxial palmar property digital artery then enters the large abaxial parietal foramen of the distal phalanx to contribute to the arcuate artery while smaller branches continue dorsally on the parietal surface of the distal phalanx (Fig 24a). Axial palmar proper digital arteries II and IV then course distally to the level of middle phalanx where the arteries turn cranially diving deep to the superficial part of the collateral ligament of the proximal interphalangeal joint and finally entering the large parietal foramen on the axial side of each distal phalanx to contribute to the arcuate artery (Fig 25a). A small dorsal common digital artery III lies on the dorsal surface in between the digital extensors (Fig 26a). Near the level of the proximal interphalangeal joint a palmar branch anastomoses with the palmar common digital artery III. At this same point the artery divides into dorsal axial proper digital artery III and dorsal axial proper digital artery IV which run down the axial surface of their respective digit to enter foramina on the proximal axial parietal surface of the distal phalanx near the extensor process.

**Figure 23.**
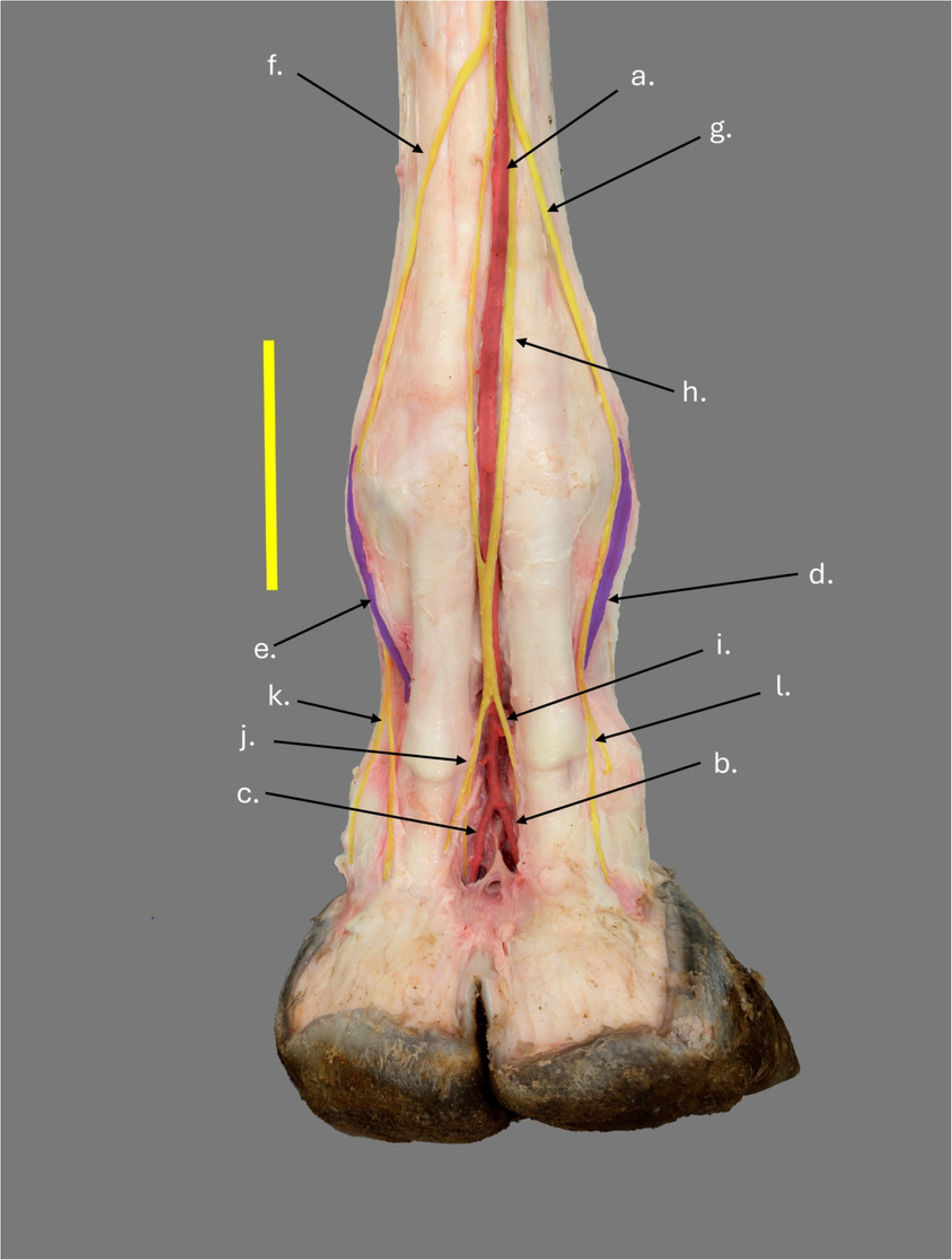
Right thoracic distal limb in palmar view of a wild Angolan giraffe. a- palmar common digital artery III. b- axial palmar proper digital artery IV. c- axial palmar proper digital artery III. d- abaxial palmar proper digital vein IV. e- abaxial palmar proper digital vein III. f-palmar common digital nerve II. g- palmar common digital nerve IV. h- palmar common digital nerve III. i- axial palmar proper digital nerve IV. j- axial palmar proper digital nerve III. k-abaxial palmar proper digital nerve III. l- abaxial palmar proper digital nerve IV. Scale bar equals 10cm.

**Figure 24.**
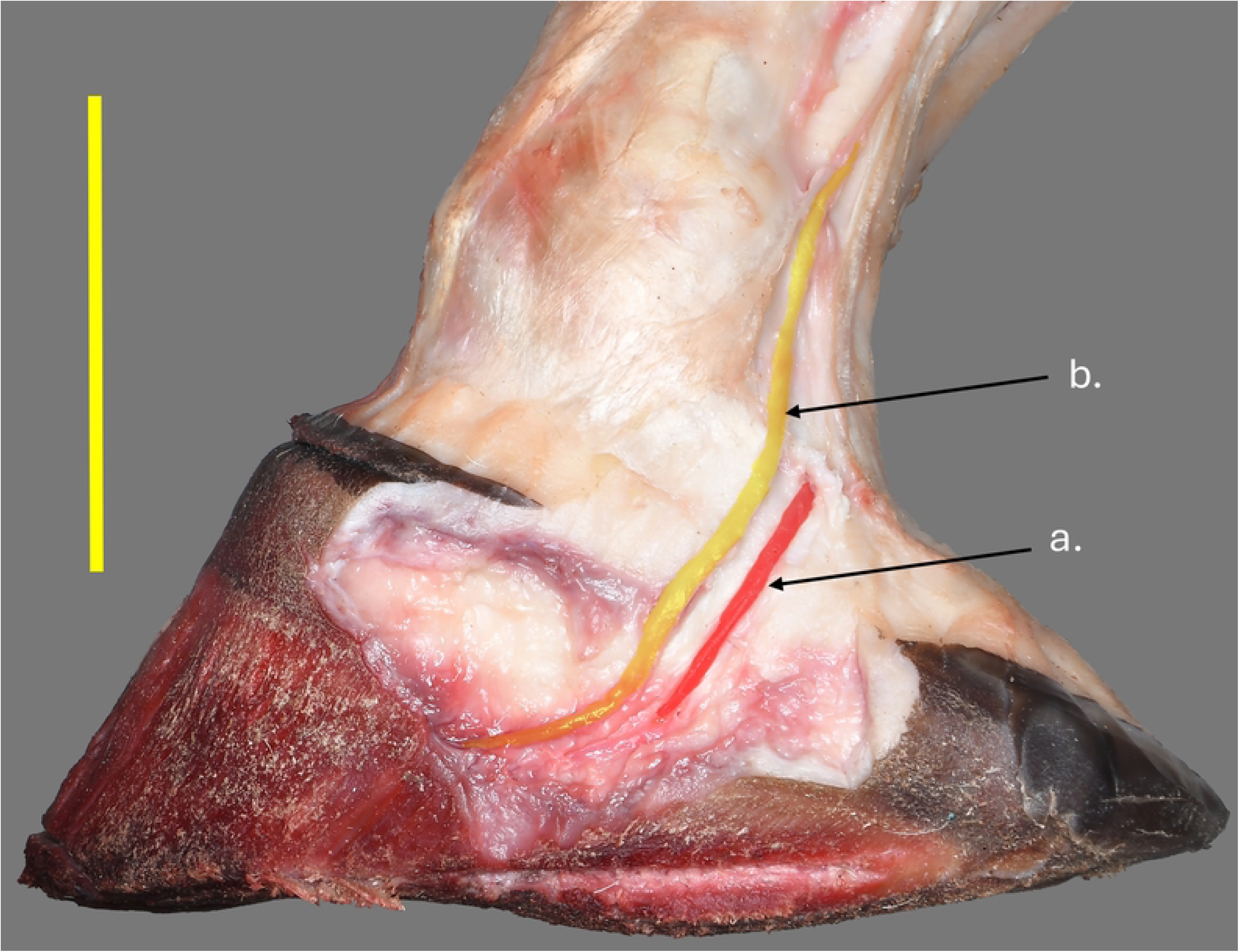
Right thoracic medial claw in abaxial view of a wild Angolan giraffe. a- abaxial palmar proper digital artery III. b- abaxial palmar proper digital nerve III. Scale bar equals 10cm.

**Figure 25.**
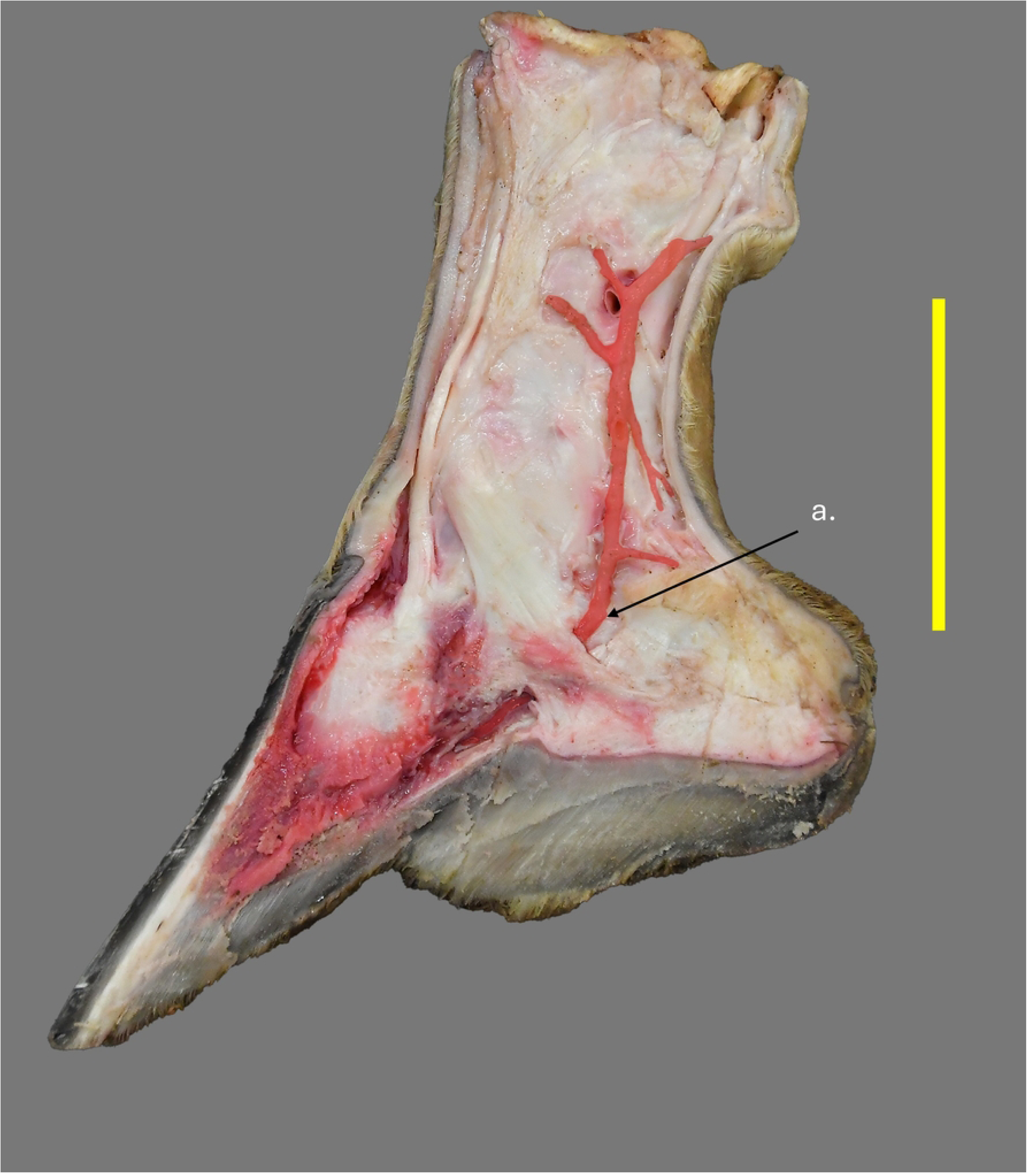
Left thoracic limb medial claw in axial view of a giraffe in human care. a- axial palmar proper digital artery III. Scale bar equals 10cm.

**Figure 26.**
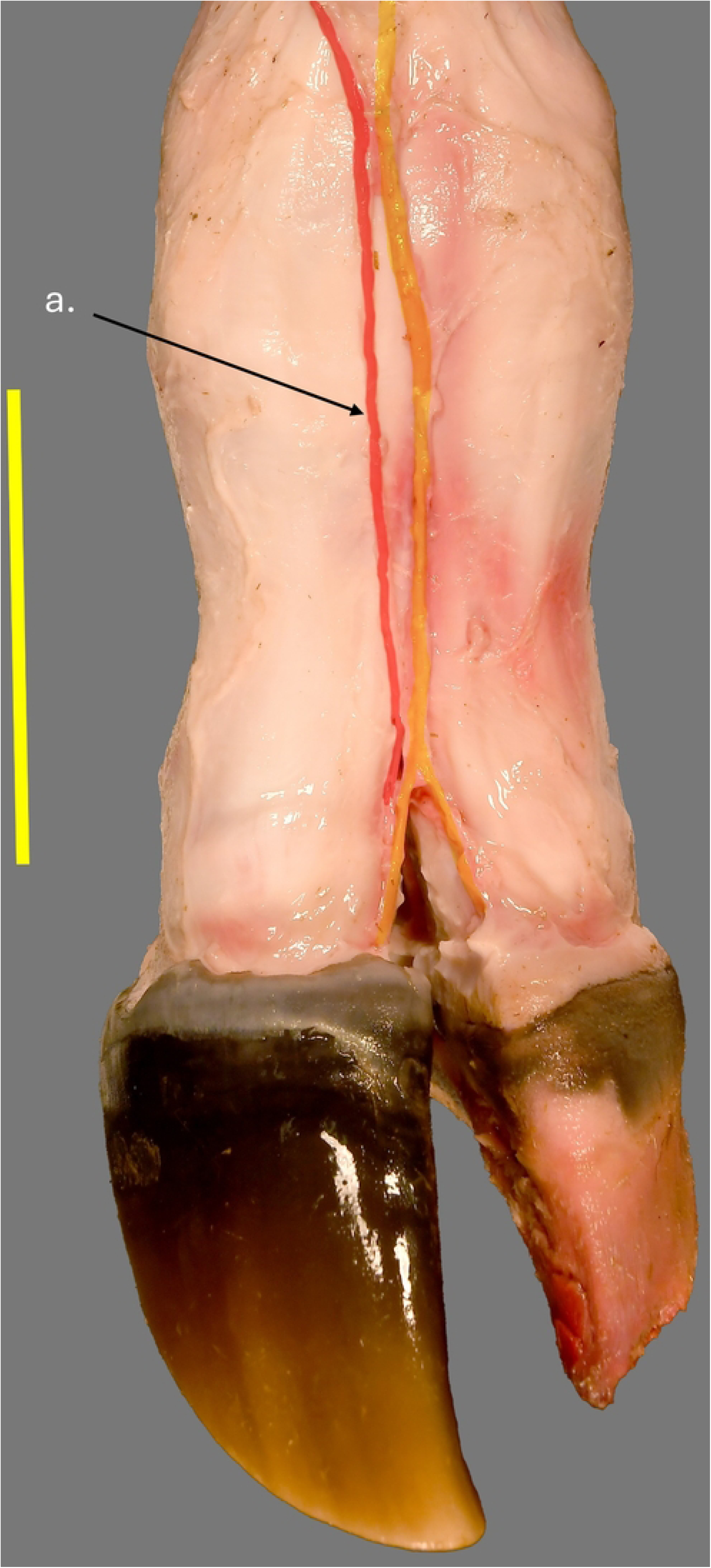
Right thoracic limb dorsal view of a giraffe in human care. a- dorsal common digital artery III. Scale bar equals 10cm.

### Arteries of the Hind Foot

Blood supply to the hind foot is primarily via dorsal metatarsal artery III which lies in a deep groove on the dorsal surface of the metatarsus deep to the digital extensor tendons (Fig. 27a). The artery runs through the dorsal intertrochlear notch at the level of the fetlock and continues in the dorsal interdigital space as dorsal common digital artery III. At a level just distal to the center of the proximal phalanx, dorsal common digital artery III makes a sharp turn running dorsal to plantar within the interdigital space (Fig. 28a). Upon reaching the level of the axial plantar surface of the digit, the artery divides into multiple branches splitting into plantar axial proper digital arteries III and IV (Fig. 29a and b). A common trunk then runs back dorsally to give the dorsal axial proper digital arteries II and IV (Fig. 27b and c) and finally several small branches supplying other regions of the foot. The proper digital arteries supply their respective digits with the plantar proper digital arteries entering the large parietal foramen in the distal phalanx and the dorsal proper digital arteries entering foramina on the dorsal surface of the distal phalanx.

**Figure 27.**
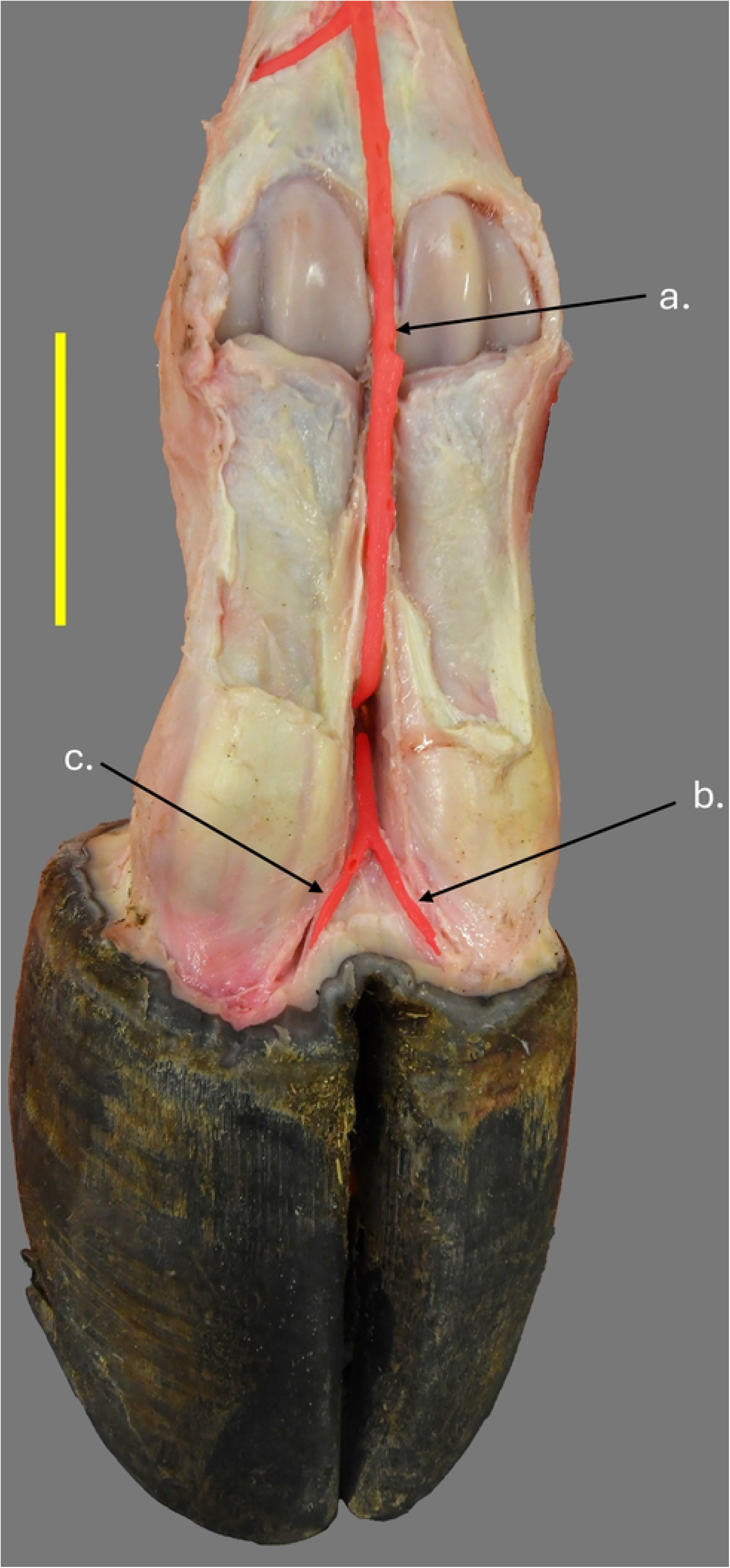
Right pelvic limb dorsal view with extensors removed of a giraffe in human care. a- dorsal metatarsal artery III. b- dorsal axial proper digital artery III. c- dorsal axial proper digital artery IV. Scale bar equals 10cm.

**Figure 28.**
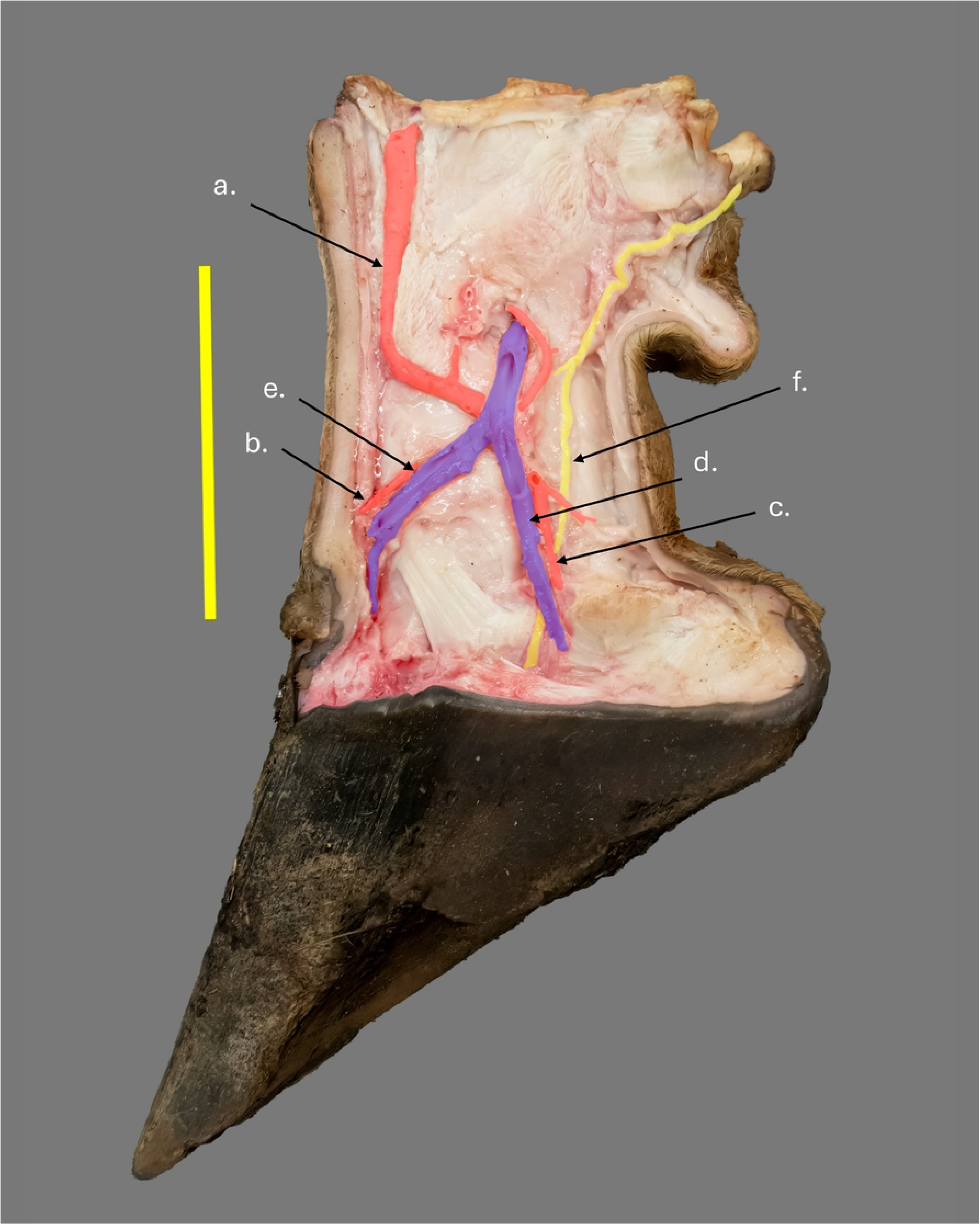
Left pelvic limb medial claw in axial view of a giraffe in human care. a- dorsal metatarsal artery III. b- dorsal axial proper digital artery III. c- axial plantar proper digital artery III. d- axial plantar proper digital vein III. e- dorsal axial proper digital vein III. f- axial plantar proper digital nerve III. Scale bar equals 10cm.

**Figure 29.**
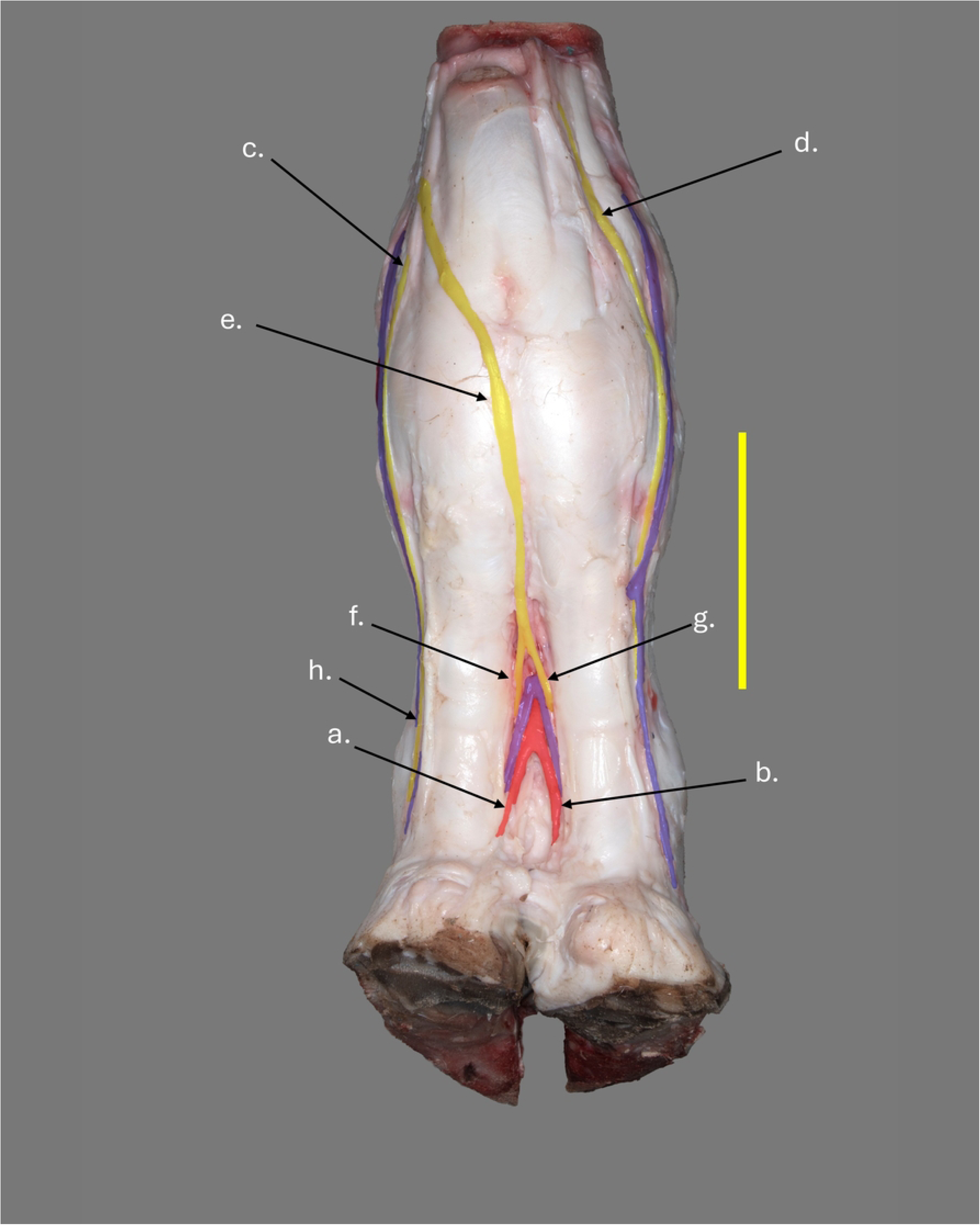
Right pelvic limb plantar view of a wild Angolan giraffe. a- axial plantar proper digital artery III. b- axial plantar proper digital artery IV. c- plantar common digital nerve II. d-plantar common digital nerve IV. e- plantar common digital nerve III. f- axial plantar proper digital nerve III. g- axial plantar proper digital nerve IV. h- abaxial plantar proper digital nerve III. Scale bar equals 10cm.

### Veins of the Front Foot

Venous drainage to the front foot is variable with various anastomoses throughout the foot. A general pattern is apparent, however. Abaxial palmar proper digital veins III and IV are relatively small and drain the abaxial side of their respective digits (Fig 23d and e). Axial palmar proper digital veins III and IV (Fig 30 and b) are much larger and come together at the level of the proximal interphalangeal joint to form palmar common digital vein III (Fig. 30c). Palmar common digital vein III runs deep in the interdigital space before giving branches which unite with the abaxial proper digital veins to form common digital vein II on the abaxial surface of digit III and common digital vein IV on the abaxial surface of digit IV (Fig 30d and e). Both common digital veins II and IV run up the palmar abaxial surfaces of their respective proximal phalanges and across the fetlock joint where they anastomose with more proximal veins deep to the interosseus (Fig. 30).

**Figure 30.**
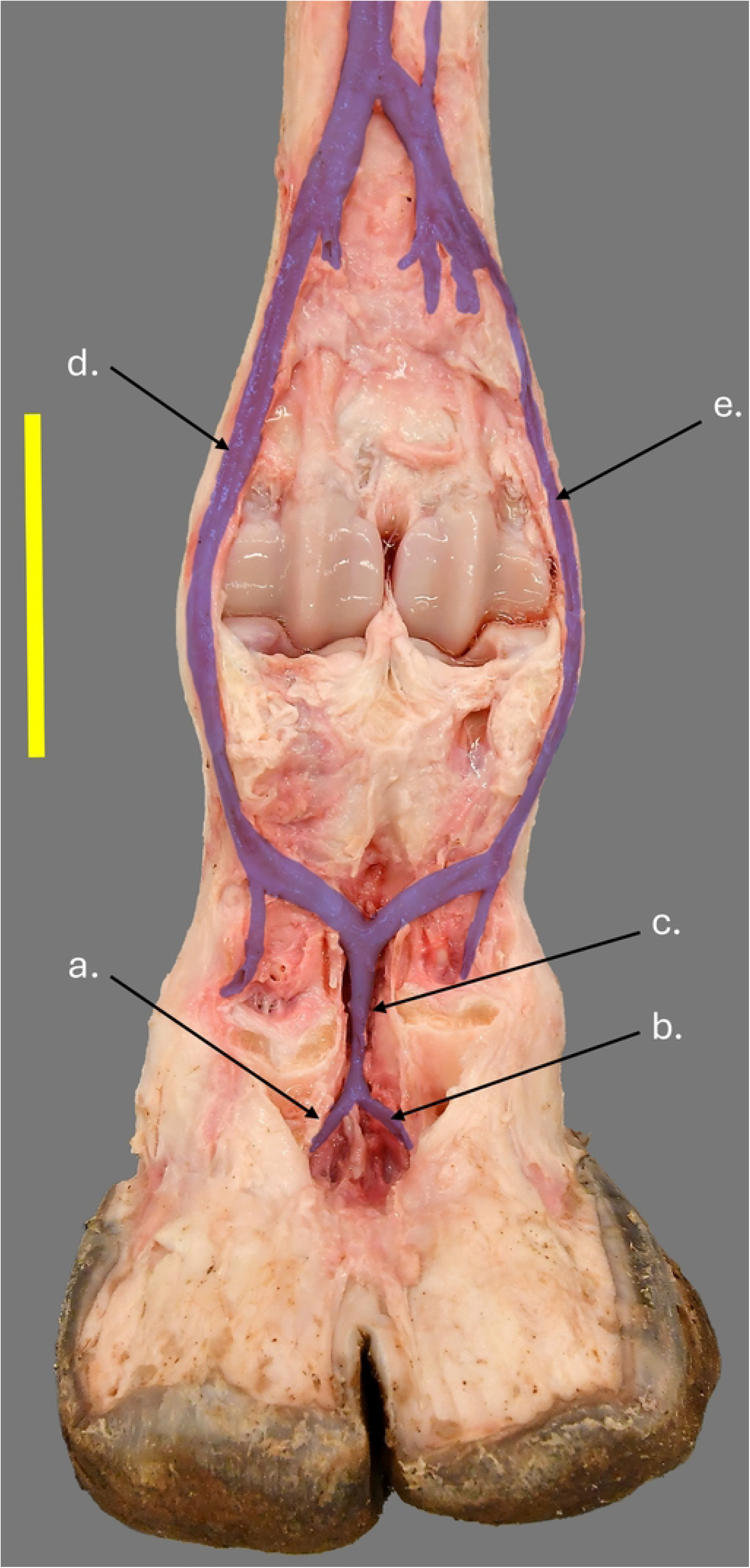
Right thoracic limb with digital flexors and interosseous removed palmar view of a giraffe in human care. a- axial palmar proper digital vein III. b- axial palmar proper digital IV. c- palmar common digital vein II. d- palmar common digital vein II. palmar common digital vein IV. Scale bar equals 10cm.

### Veins of the Hind Foot

Venous drainage of the hind foot is also variable, especially in the region of the digits. There are numerous anastomoses as the veins course proximally. Some branches do, however, seem to be consistent. Dorsal axial proper digital veins III and IV (Fig 28e) and plantar axial proper digital veins III and IV (Fig 28d and Fig 31b and c) parallel the course of their respective arteries and unite in the interdigital space to form common plantar digital vein III at the level of the proximal interphalangeal joint (Fig. 31d). Plantar abaxial proper digital vein III was found on all specimens (Fig 31a) but at least one specimen lacked plantar abaxial proper digital vein IV (Fig. 31). When present, both lie close to the digital flexors on the plantar surface of the foot. Plantar common digital vein III in the center of the interdigital space divides as it moves proximally giving branches which unite with the plantar abaxial proper digital branches to form plantar common digital vein II (Fig 31e) on the abaxial side of digit III and plantar common digital vein IV (Fig 31f) on the abaxial side of digit IV. Plantar common digital veins II and IV then run on the abaxial side of the fetlock joint just under the skin before moving deep to the interosseus on the caudal surface of the metatarsus (Fig 31).

**Figure 31.**
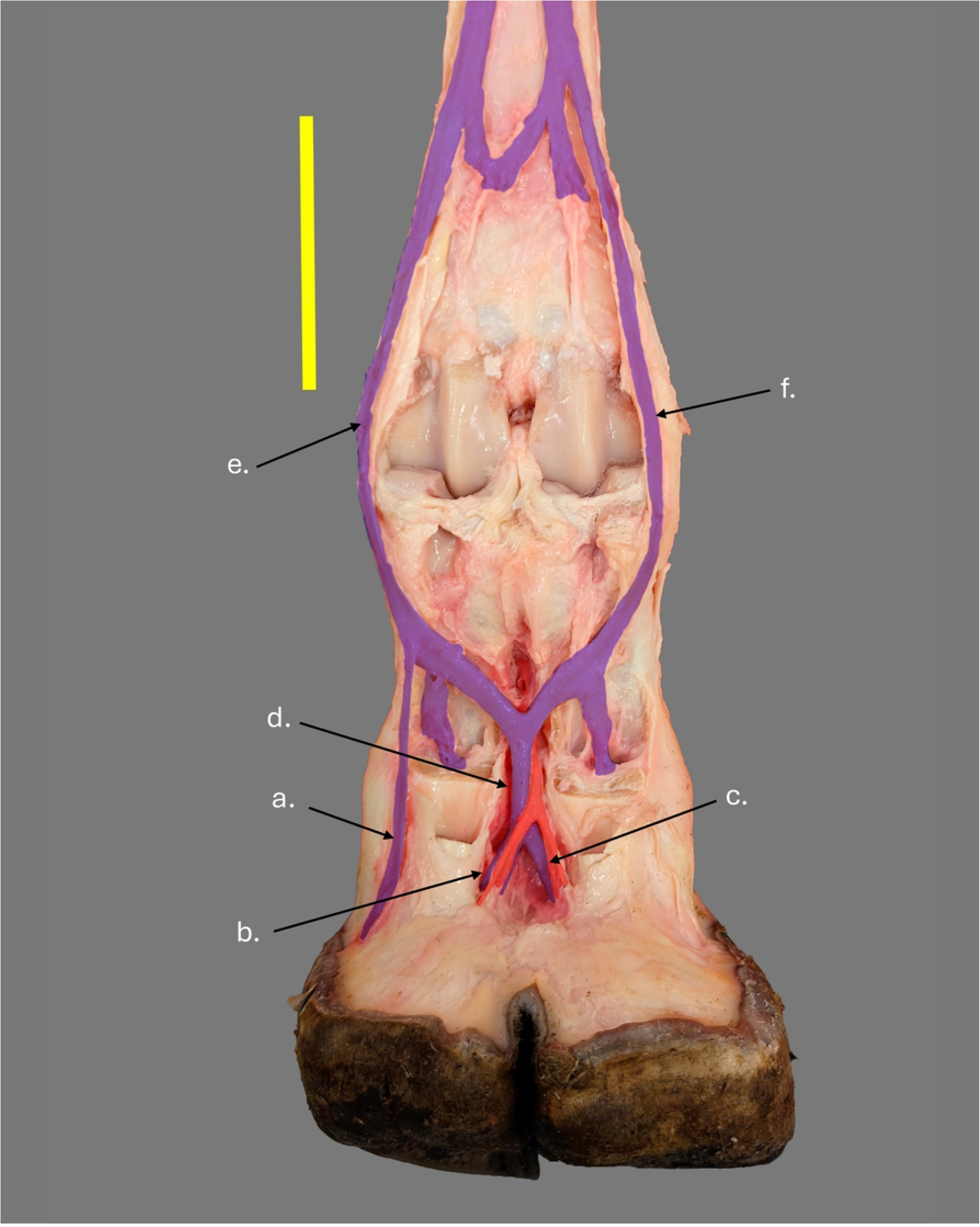
Right pelvic limb with digital flexors and interosseous removed plantar view of a giraffe in human care. a- abaxial plantar proper digital vein III. b- axial plantar proper digital vein III. c- axial plantar proper digital vein IV. d- plantar common digital vein III. e- plantar common digital vein II. f- plantar common digital vein IV. Scale bar equals 10cm.

### Nerves of the Front Foot

Approximately 15 cm above the fetlock, where the limbs were cut, the median nerve is already divided into palmar common digital nerve II (Fig 23f), palmar common digital nerve III (Fig 23h) and palmar common digital nerve IV (Fig 23g). Palmar common digital nerve II runs down the abaxial side of the fetlock and onto the abaxial palmar surface of digit III. At this point it is abaxial palmar proper digital nerve III (Fig 23k) and follows the abaxial palmar proper digital vessels of digit III into the foot (Fig 24b). Palmar common digital nerve III is separated into two branches and lies on either side of palmar common digital artery III at the level of the fetlock (Fig 23h). After crossing the fetlock, in some individuals the branches unite to form a single palmar common digital nerve III. In these individuals, the united nerve then runs in the interdigital space distal to the level of the proximal digital annular ligaments where it divides into axial palmar digital nerve III and axial palmar proper digital nerve IV (Fig 23i and j). The proper palmar axial nerves then run down the palmar axial surface of their respective digits and follow the proper palmar axial vessels into the foot. Palmar common digital nerve IV can be seen crossing the surface of the superficial digital flexor from the medial aspect of the distal metacarpal to the lateral surface (Fig 23g). Once palmar common digital nerve IV reaches the abaxial surface of digit IV and crosses the fetlock joint, it is referred to as abaxial palmar proper digital nerve IV (Fig 23l). At this point the nerve follows the abaxial palmar proper digital vessels of digit IV into the foot.

### Nerves of the Hind Foot

Approximately 20 cm above the fetlock joint, the medial plantar nerve gives off plantar common digital nerve IV which runs over the plantar surface of the superficial digital flexor to course down the lateral side of the distal metatarsus (Fig 29d). After crossing the fetlock joint, the nerve continues as abaxial plantar proper digital nerve IV, running with the plantar abaxial proper digital vessels of digit IV into the foot. A few centimeters distal to the branching of the plantar common digital nerve IV, the medial plantar nerve terminates by splitting into plantar common digital nerve III (Fig 29e) and plantar common digital nerve II (Fig 29c). Plantar common digital nerve II then continues across the medial side of the metatarsus until it crosses the fetlock, where it continues as the abaxial plantar proper digital nerve II (Fig 29h), running with the plantar abaxial proper digital vessels of digit II into the foot. The plantar common digital nerve III runs across the plantar surface of the medial palmar annular ligament and reaches the interdigital space at the level of the proximal sesamoids (Fig 29e). The nerve then runs in the interdigital space superficial to plantar common digital vein III and the plantar common digital artery III. Just proximal to the level of the proximal digital annular ligament, plantar common digital nerve III divides into axial plantar proper digital nerve III (Fig 29f) and axial plantar proper digital nerve IV (Fig 29g). Each axial palmar proper digital nerve then runs down the axial surface of its respective digit in the interdigital space before continuing into the foot with their respective vessels.

### Hoof Capsule

The hoof capsules of the front feet are significantly larger than those of the hind feet (Fig 32). In both the front and hind feet of wild giraffe, the lateral hoof capsule is larger than the medial hoof capsule (Fig 32). Asymmetry was commonly observed in both front feet, particularly in toe-tip morphology, with one toe appearing round and the other more pointed (Fig 32). Variations in overall hoof capsule shape and length were also noted. Thickness of the hoof capsule varies from one region to another. The thickest area of the hoof capsule is found in the sole (Figs 33a and 34c). The abaxial hoof wall is the next thickest (Fig 34b) followed by the axial hoof wall which is both thin and short (Fig 34a). The distal surface of the hoof capsule can be divided into a broad flat heel region and an axially concave toe region (Fig 32). The hoof capsule is tallest at the toe and tapers steeply to the heel where the hoof wall is very short (Fig 4B and D). Table 1 shows the hoof capsule measurements for the seven specimens used in this study. When removing the distal surface of the hoof capsule from wild feet, it was common for the hoof horn to separate under the flat heel portion of the distal surface (Fig 35). The heel region of the sole exhibited mechanical properties which lead us to believe that the horn in that region is more malleable than that of the rest of the sole. This was expressed in the hardness of the two portions of the distal surface with the toe region being much harder than the heel region.

**Figure 32.**
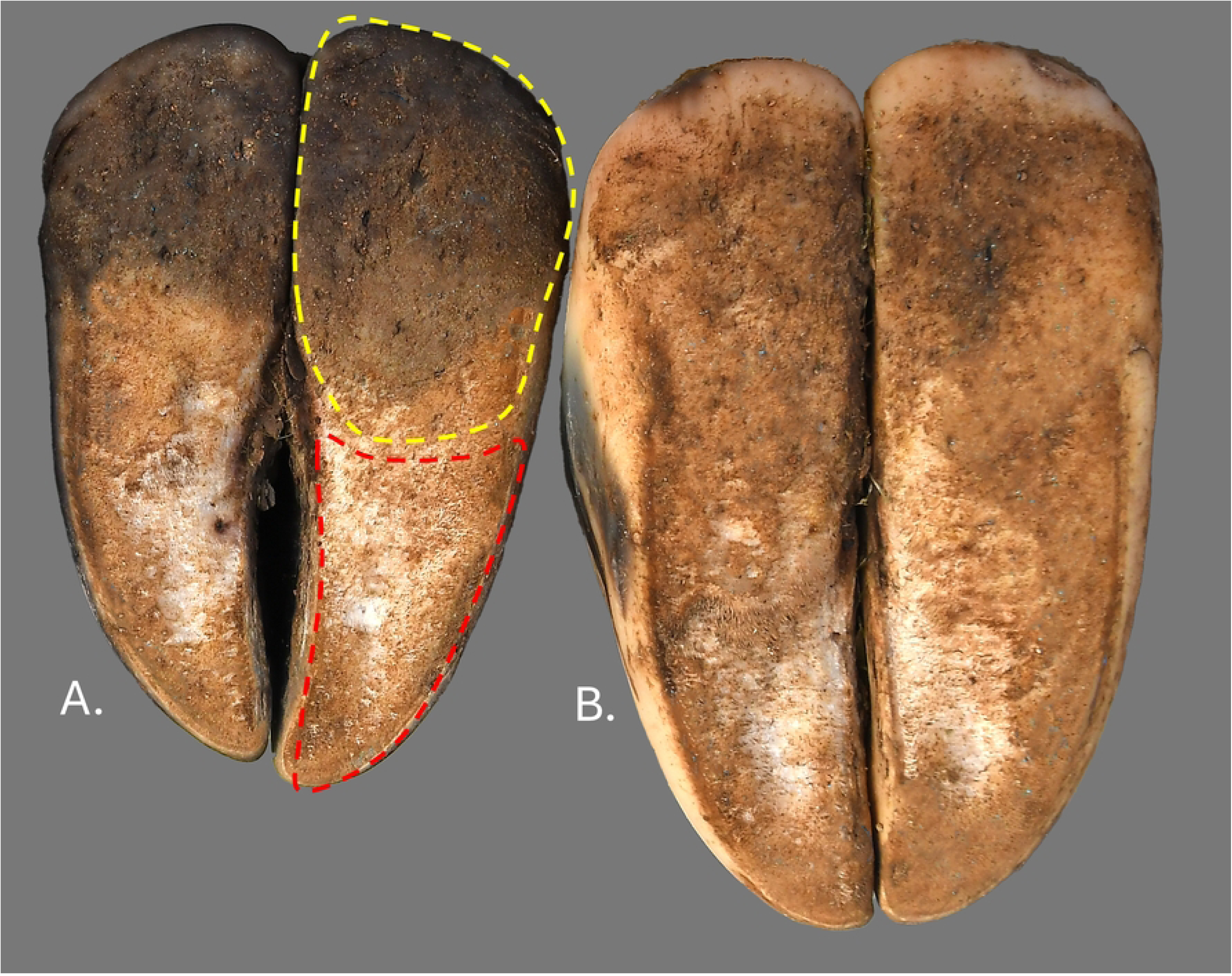
Right limbs in distal view from a single individual of a wild Angolan giraffe. (A) Pelvic limb. (B) Thoracic limb. The yellow dotted outline shows the heel region of the right lateral digit of the hind foot. The red dotted outline shows the toe region of the right lateral digit of the hind foot.

**Figure 33.**
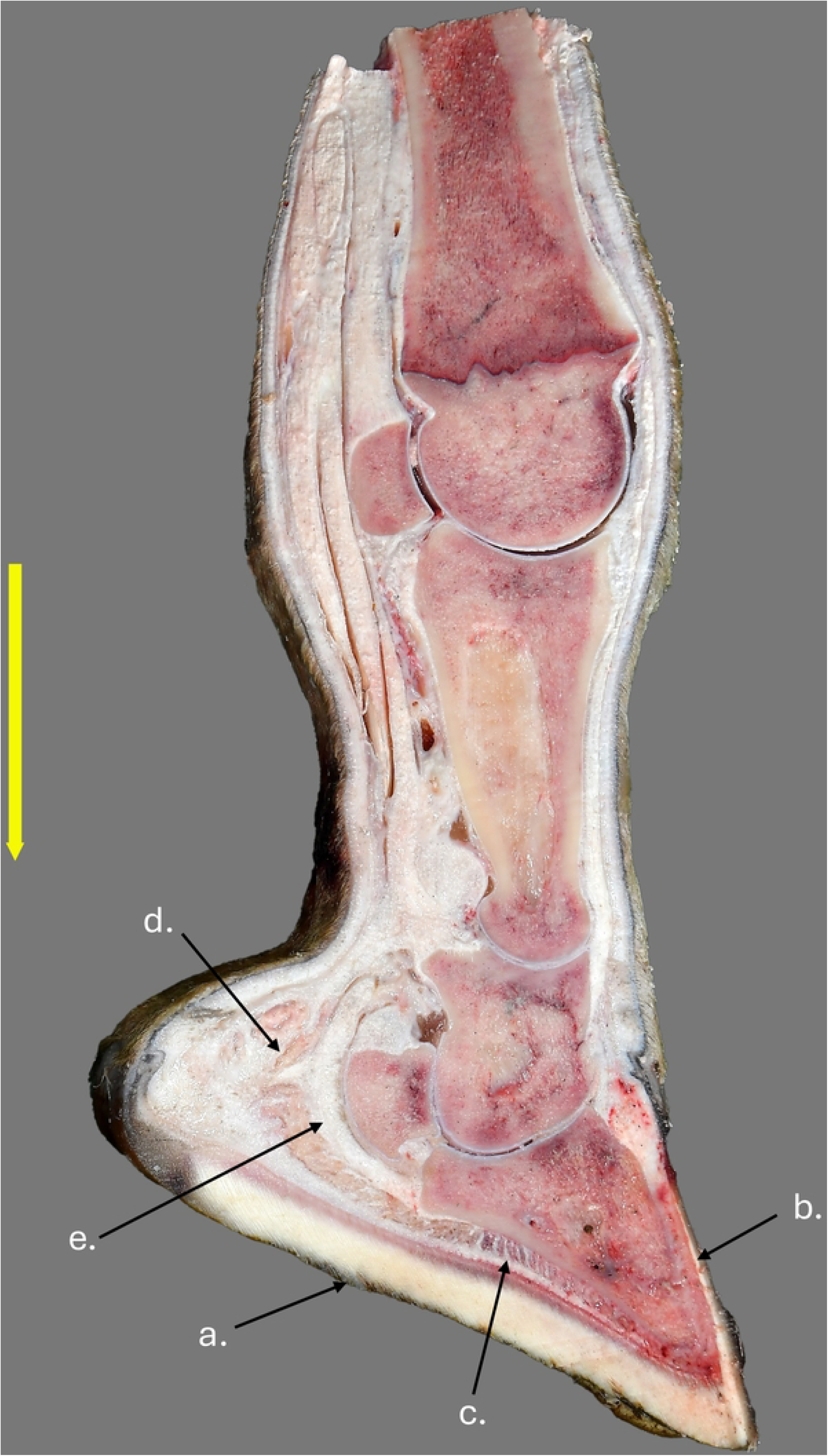
Mid-sagittal section through the left thoracic limb medial claw of a wild Angolan giraffe. a- sole. b- hoof wall. c- digital cushion beneath distal phalanx. d- heel bulb portion of the digital cushion. e- deep digital flexor. Scale bar equals 10cm.

**Figure 34.**
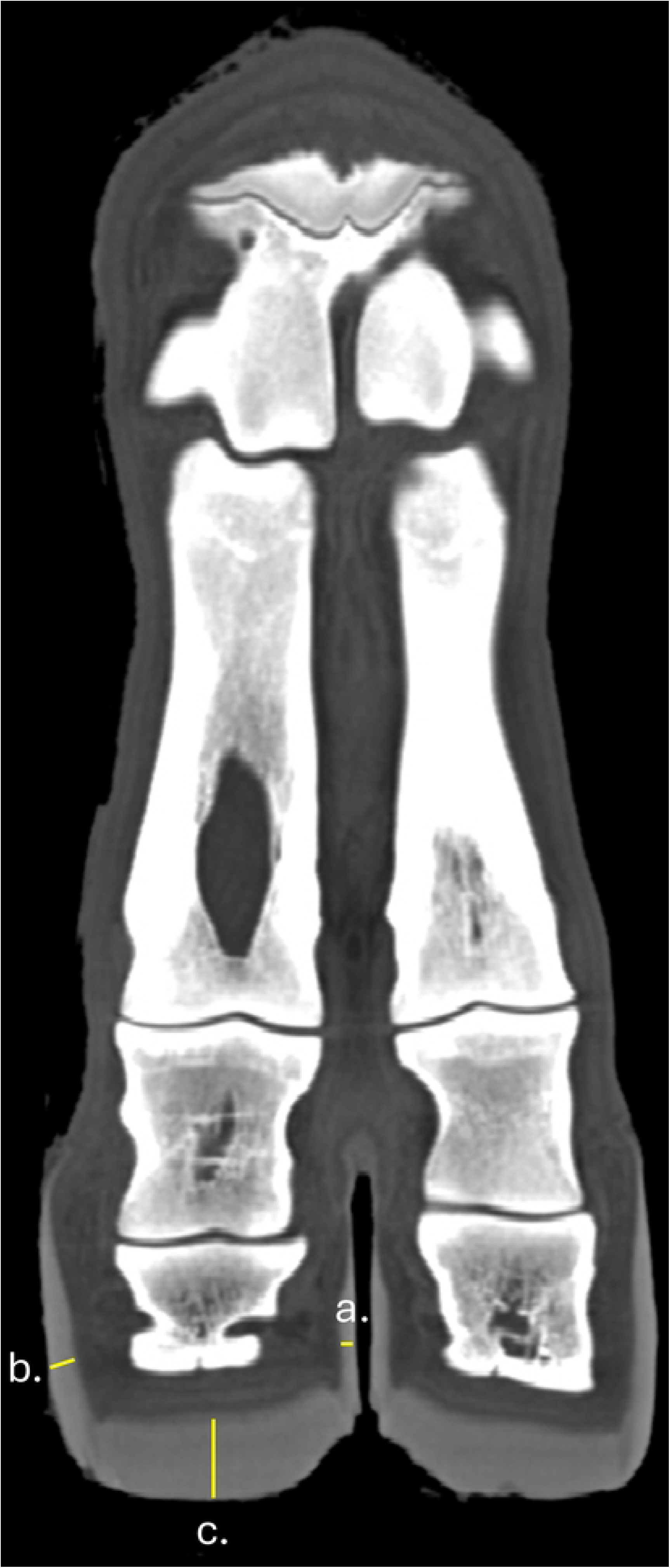
CT left thoracic coronal section of a wild Angolan giraffe. Yellow lines indicate the thickness of the hoof capsule at a- axial hoof wall, b- abaxial hoof wall, and c- sole.

**Figure 35.**
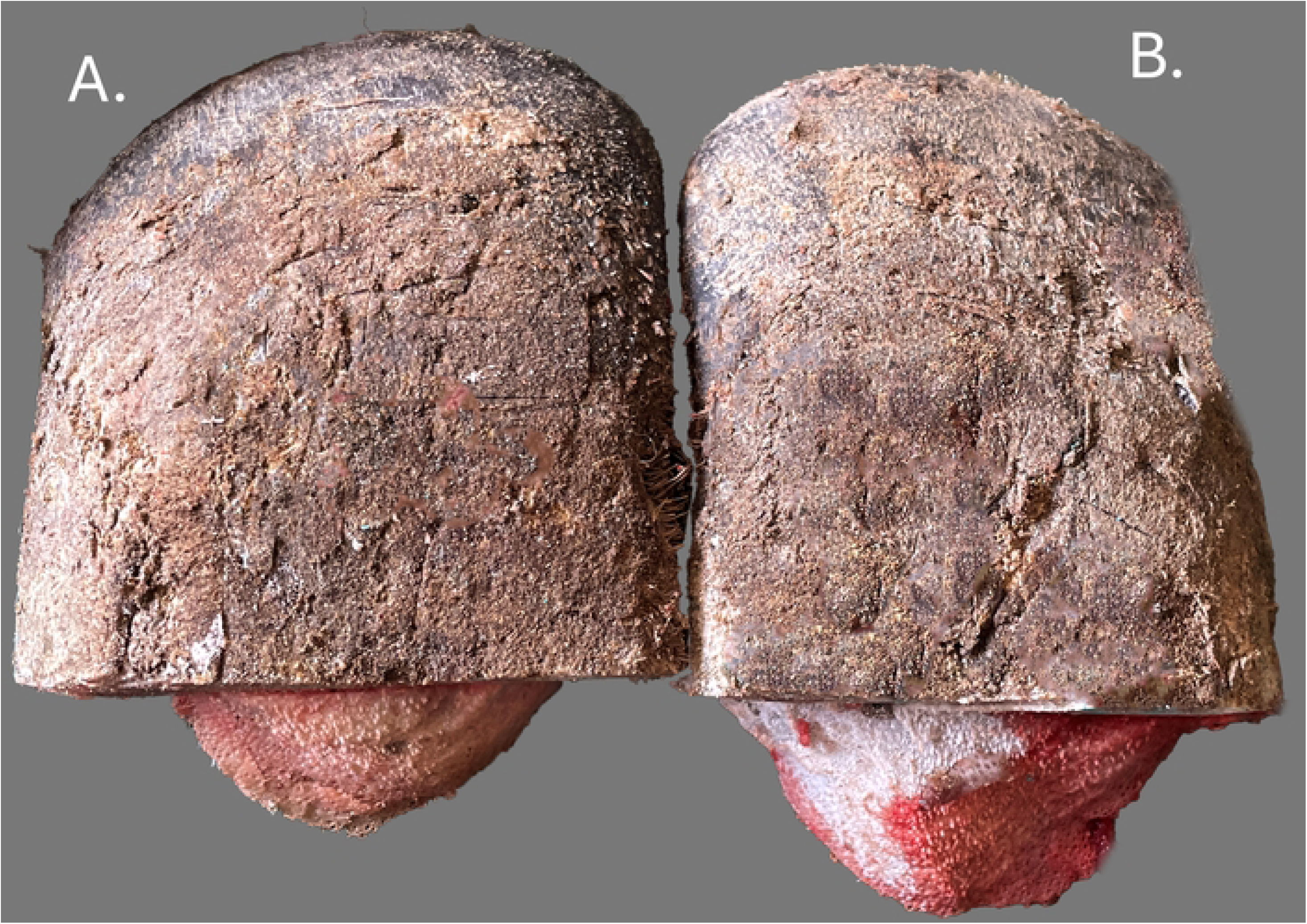
Left thoracic heel region of wild Angolan giraffe hoof showing a pattern of differential separation of hoof sole layers. (A) Lateral heel. (B) Medial heel.

**Table 1.**
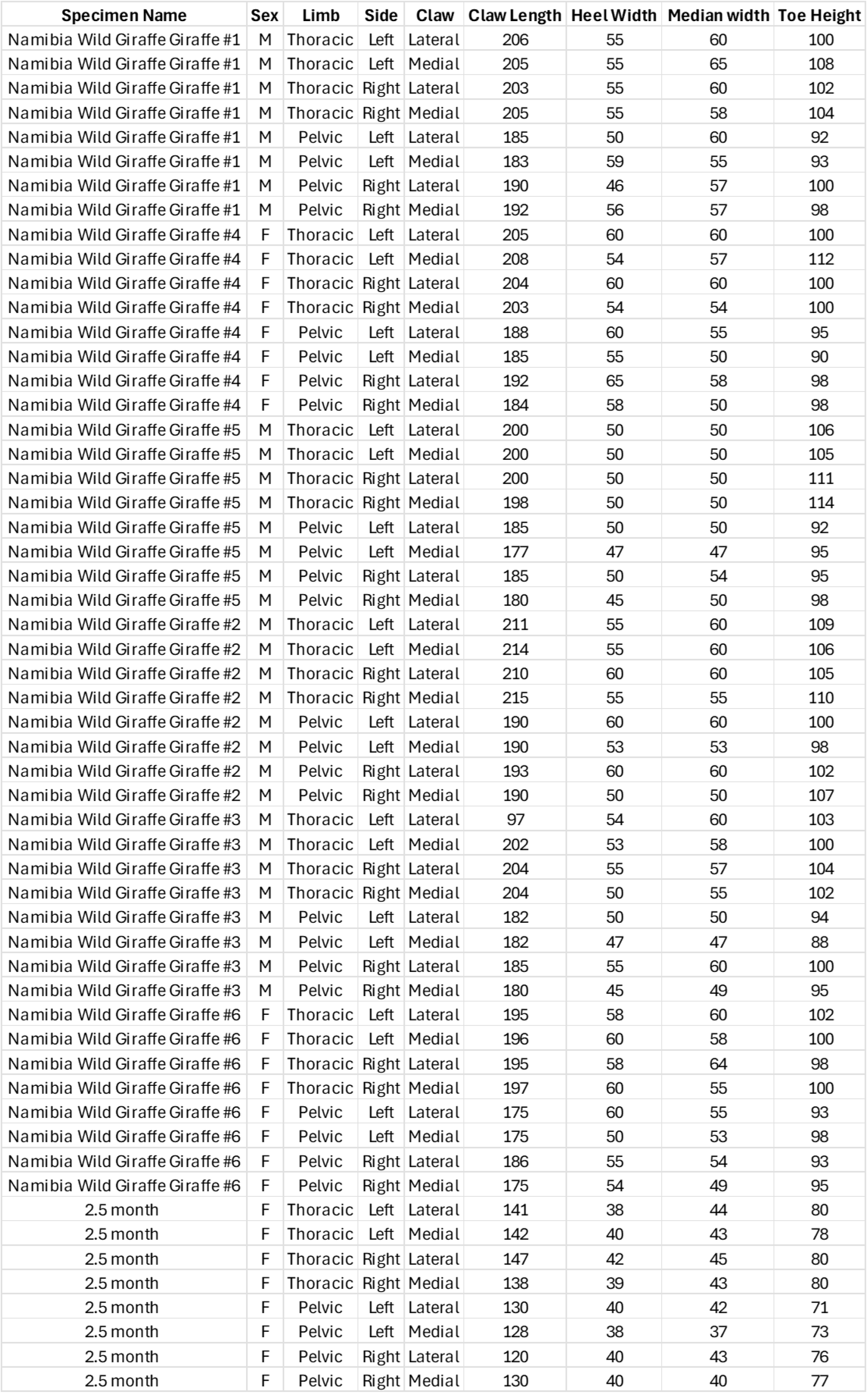
Foot measurements (in millimeters) for wild Angolan giraffe (*G. giraffa angolensis*) and northern giraffe (*G. camelopardalis*) under human care feet used in the study.

| Specimen Name | Sex | Limb | Side | Claw | Claw Length | Heel Width | Median width | Toe Height |
| --- | --- | --- | --- | --- | --- | --- | --- | --- |
| Namibia Wild Giraffe Giraffe #1 | M | Thoracic | Left | Lateral | 206 | 55 | 60 | 100 |
| Namibia Wild Giraffe Giraffe #1 | M | Thoracic | Left | Medial | 205 | 55 | 65 | 108 |
| Namibia Wild Giraffe Giraffe #1 | M | Thoracic | Right | Lateral | 203 | 55 | 60 | 102 |
| Namibia Wild Giraffe Giraffe #1 | M | Thoracic | Right | Medial | 205 | 55 | 58 | 104 |
| Namibia Wild Giraffe Giraffe #1 | M | Pelvic | Left | Lateral | 185 | 50 | 60 | 92 |
| Namibia Wild Giraffe Giraffe #1 | M | Pelvic | Left | Medial | 183 | 59 | 55 | 93 |
| Namibia Wild Giraffe Giraffe #1 | M | Pelvic | Right | Lateral | 190 | 46 | 57 | 100 |
| Namibia Wild Giraffe Giraffe #1 | M | Pelvic | Right | Medial | 192 | 56 | 57 | 98 |
| Namibia Wild Giraffe Giraffe #4 | F | Thoracic | Left | Lateral | 205 | 60 | 60 | 100 |
| Namibia Wild Giraffe Giraffe #4 | F | Thoracic | Left | Medial | 208 | 54 | 57 | 112 |
| Namibia Wild Giraffe Giraffe #4 | F | Thoracic | Right | Lateral | 204 | 60 | 60 | 100 |
| Namibia Wild Giraffe Giraffe #4 | F | Thoracic | Right | Medial | 203 | 54 | 54 | 100 |
| Namibia Wild Giraffe Giraffe #4 | F | Pelvic | Left | Lateral | 188 | 60 | 55 | 95 |
| Namibia Wild Giraffe Giraffe #4 | F | Pelvic | Left | Medial | 185 | 55 | 50 | 90 |
| Namibia Wild Giraffe Giraffe #4 | F | Pelvic | Right | Lateral | 192 | 65 | 58 | 98 |
| Namibia Wild Giraffe Giraffe #4 | F | Pelvic | Right | Medial | 184 | 58 | 50 | 98 |
| Namibia Wild Giraffe Giraffe #5 | M | Thoracic | Left | Lateral | 200 | 50 | 50 | 106 |
| Namibia Wild Giraffe Giraffe #5 | M | Thoracic | Left | Medial | 200 | 50 | 50 | 105 |
| Namibia Wild Giraffe Giraffe #5 | M | Thoracic | Right | Lateral | 200 | 50 | 50 | 111 |
| Namibia Wild Giraffe Giraffe #5 | M | Thoracic | Right | Medial | 198 | 50 | 50 | 114 |
| Namibia Wild Giraffe Giraffe #5 | M | Pelvic | Left | Lateral | 185 | 50 | 50 | 92 |
| Namibia Wild Giraffe Giraffe #5 | M | Pelvic | Left | Medial | 177 | 47 | 47 | 95 |
| Namibia Wild Giraffe Giraffe #5 | M | Pelvic | Right | Lateral | 185 | 50 | 54 | 95 |
| Namibia Wild Giraffe Giraffe #5 | M | Pelvic | Right | Medial | 180 | 45 | 50 | 98 |
| Namibia Wild Giraffe Giraffe #2 | M | Thoracic | Left | Lateral | 211 | 55 | 60 | 109 |
| Namibia Wild Giraffe Giraffe #2 | M | Thoracic | Left | Medial | 214 | 55 | 60 | 106 |
| Namibia Wild Giraffe Giraffe #2 | M | Thoracic | Right | Lateral | 210 | 60 | 60 | 105 |
| Namibia Wild Giraffe Giraffe #2 | M | Thoracic | Right | Medial | 215 | 55 | 55 | 110 |
| Namibia Wild Giraffe Giraffe #2 | M | Pelvic | Left | Lateral | 190 | 60 | 60 | 100 |
| Namibia Wild Giraffe Giraffe #2 | M | Pelvic | Left | Medial | 190 | 53 | 53 | 98 |
| Namibia Wild Giraffe Giraffe #2 | M | Pelvic | Right | Lateral | 193 | 60 | 60 | 102 |
| Namibia Wild Giraffe Giraffe #2 | M | Pelvic | Right | Medial | 190 | 50 | 50 | 107 |
| Namibia Wild Giraffe Giraffe #3 | M | Thoracic | Left | Lateral | 97 | 54 | 60 | 103 |
| Namibia Wild Giraffe Giraffe #3 | M | Thoracic | Left | Medial | 202 | 53 | 58 | 100 |
| Namibia Wild Giraffe Giraffe #3 | M | Thoracic | Right | Lateral | 204 | 55 | 57 | 104 |
| Namibia Wild Giraffe Giraffe #3 | M | Thoracic | Right | Medial | 204 | 50 | 55 | 102 |
| Namibia Wild Giraffe Giraffe #3 | M | Pelvic | Left | Lateral | 182 | 50 | 50 | 94 |
| Namibia Wild Giraffe Giraffe #3 | M | Pelvic | Left | Medial | 182 | 47 | 47 | 88 |
| Namibia Wild Giraffe Giraffe #3 | M | Pelvic | Right | Lateral | 185 | 55 | 60 | 100 |
| Namibia Wild Giraffe Giraffe #3 | M | Pelvic | Right | Medial | 180 | 45 | 49 | 95 |
| Namibia Wild Giraffe Giraffe #6 | F | Thoracic | Left | Lateral | 195 | 58 | 60 | 102 |
| Namibia Wild Giraffe Giraffe #6 | F | Thoracic | Left | Medial | 196 | 60 | 58 | 100 |
| Namibia Wild Giraffe Giraffe #6 | F | Thoracic | Right | Lateral | 195 | 58 | 64 | 98 |
| Namibia Wild Giraffe Giraffe #6 | F | Thoracic | Right | Medial | 197 | 60 | 55 | 100 |
| Namibia Wild Giraffe Giraffe #6 | F | Pelvic | Left | Lateral | 175 | 60 | 55 | 93 |
| Namibia Wild Giraffe Giraffe #6 | F | Pelvic | Left | Medial | 175 | 50 | 53 | 98 |
| Namibia Wild Giraffe Giraffe #6 | F | Pelvic | Right | Lateral | 186 | 55 | 54 | 93 |
| Namibia Wild Giraffe Giraffe #6 | F | Pelvic | Right | Medial | 175 | 54 | 49 | 95 |
| 2.5 month | F | Thoracic | Left | Lateral | 141 | 38 | 44 | 80 |
| 2.5 month | F | Thoracic | Left | Medial | 142 | 40 | 43 | 78 |
| 2.5 month | F | Thoracic | Right | Lateral | 147 | 42 | 45 | 80 |
| 2.5 month | F | Thoracic | Right | Medial | 138 | 39 | 43 | 80 |
| 2.5 month | F | Pelvic | Left | Lateral | 130 | 40 | 42 | 71 |
| 2.5 month | F | Pelvic | Left | Medial | 128 | 38 | 37 | 73 |
| 2.5 month | F | Pelvic | Right | Lateral | 120 | 40 | 43 | 76 |
| 2.5 month | F | Pelvic | Right | Medial | 130 | 40 | 40 | 77 |

### Digital Cushion and Heel Bulbs

The hypodermis under the distal phalanx constitutes the digital cushion and consists of multiple narrow compartments running axial to abaxial filled with adipose tissue and separated by corresponding lemellae of dense connective tissue (Figs 33c and 36). Behind the distal phalanx, the digital cushion expands to form a large portion of the heel bulb which consists of two large fat bodies separated by a strong dense connective tissue lamina running in a dorsal to palmar/plantar plane (Fig 36). The fat bodies are further encapsulated in a dense fibrous capsule that interdigitates with fibers of the distal digital annular ligament as well as the distal surface of the tendon of insertion of the deep digital flexor (Fig 33d).

**Figure 36.**
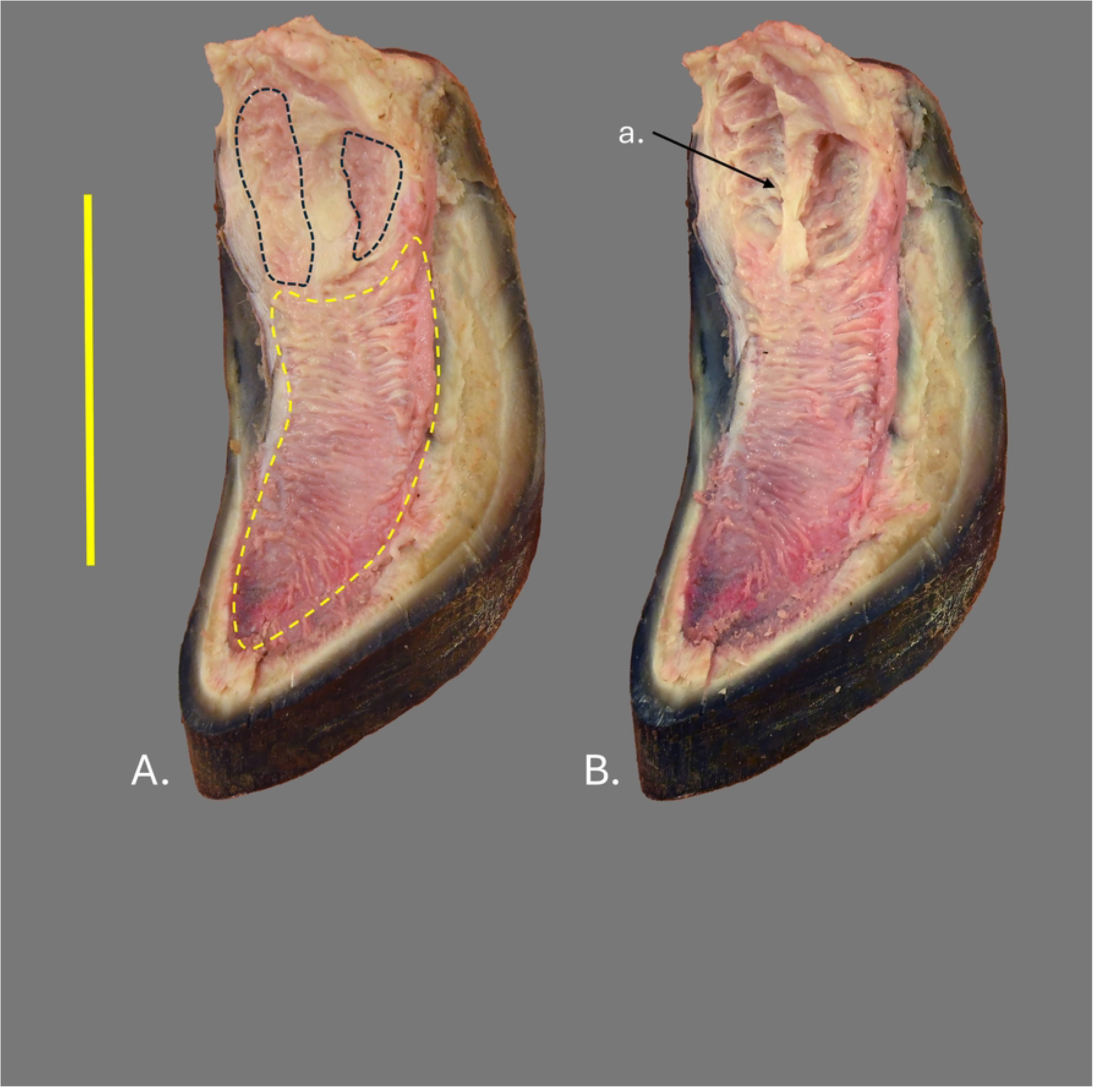
Isolated digital cushion and sole from the medial claw of the right pelvic limb in proximal view of a giraffe in human care. (A) With the adipose component of the heel bulbs intact. (B) With the adipose component of the heel bulbs removed. a- dense connective tissue septum dividing the adipose portion of the digital cushion in the heel bulb into axial and abaxial components. The yellow dotted outline is the portion of the digital cushion directly under the distal phalanx. The black dotted outlines are the adipose components of the digital cushion in the heel bulb. Scale bar equals 10cm.

After segmentation of the distal phalanges and adipose portion of the heel bulbs (Fig 37), relative ages for the six wild Angolan giraffe (*G. giraffa angolensis*) were determined based on patterns of epiphyseal suture closure. The ratio of adipose heel bulb volume to distal phalanx volume can be seen in Figure 38 with specimens arranged from youngest to oldest based on epiphyseal suture closure and actual measurements of distal phalanx and adipose heel bulb volume can be seen in Table 2. Several patterns are clearly visible. Both the volume of the distal phalanx and the adipose heel bulb are greater in the front feet than the hind feet. Further, within each foot, the lateral distal phalanx and heel bulb are larger than the medial distal phalanx and heel bulb on both the front and hind feet. Also, there appears to be a pattern of decreasing adipose heel bulb volume when compared to distal phalanx volume with increased age. The lone exception being the right rear medial claw which shows little change between the 2.5 month old northern giraffe (*G. camelopardalis*) and the youngest wild Angolan giraffe (*G. giraffa angolensis*) specimens.

**Figure 37.**
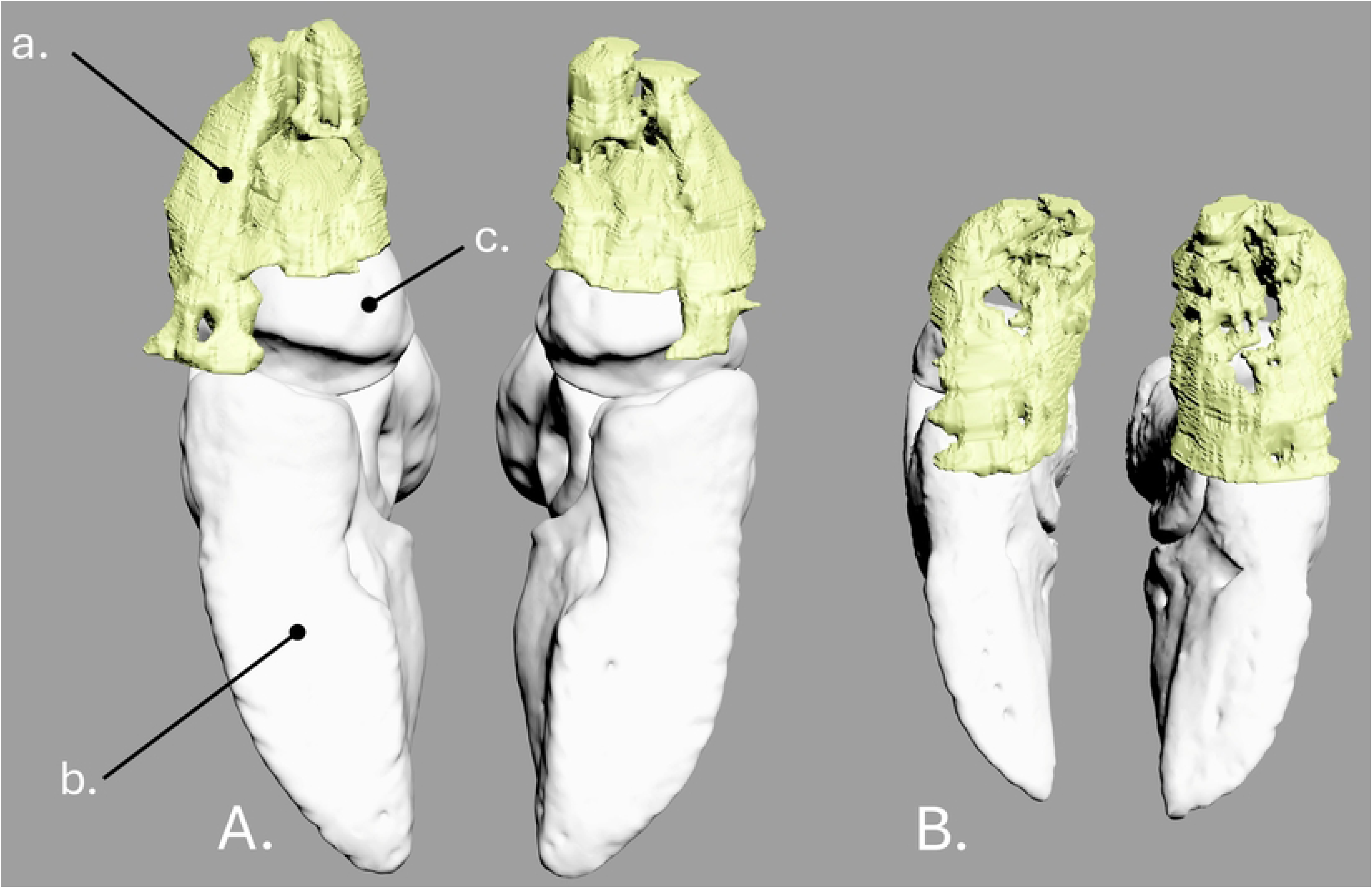
3-D reconstruction of the right thoracic limb of (A) adult wild Angolan giraffe (B) 2.5 month old human care giraffe in distal view. a- adipose tissue of the heel bulb. b-distal phalanx. c- distal sesamoid (navicular bone).

**Figure 38.**
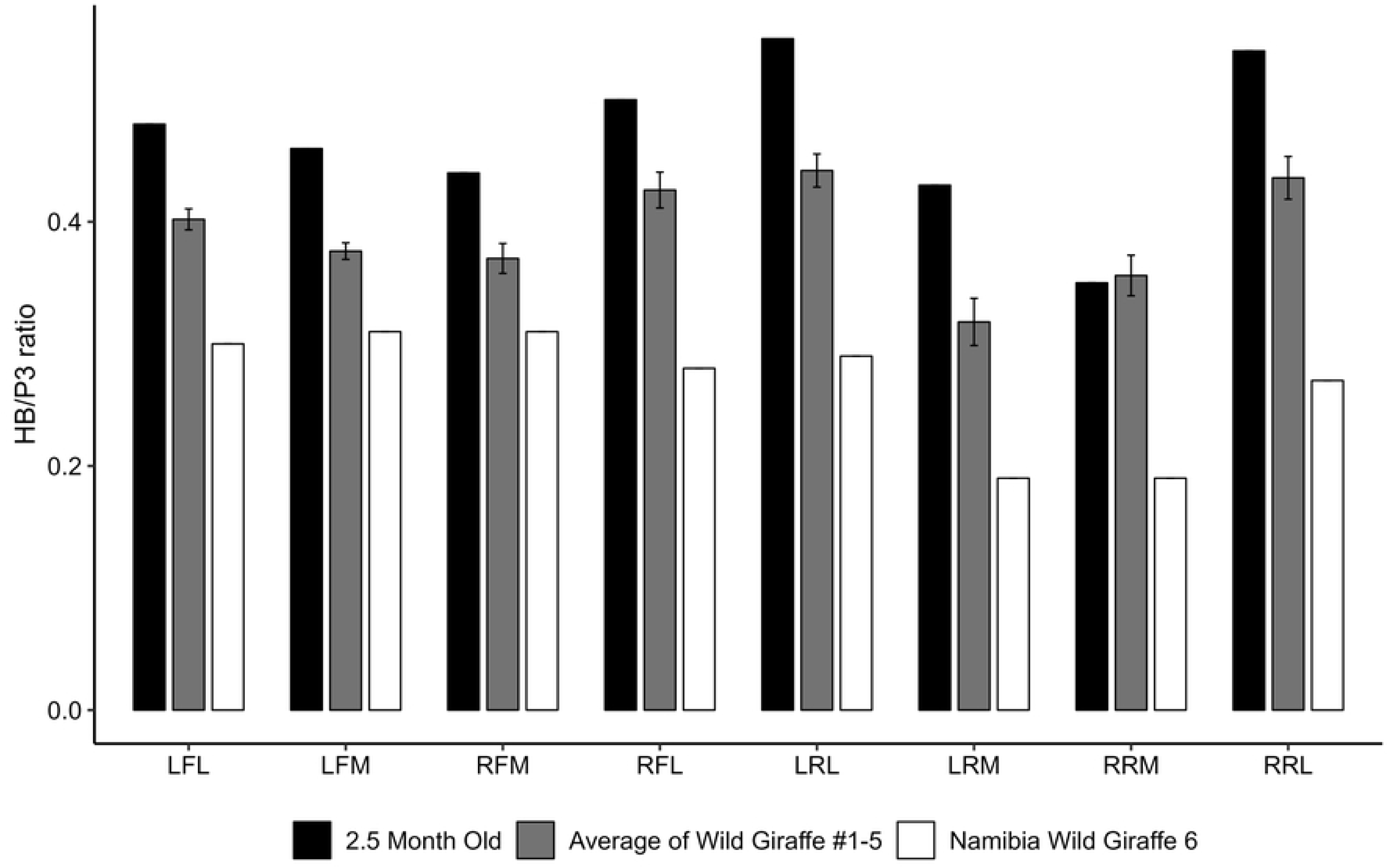
Ratio of volume of adipose heel bulb to the volume of distal phalanx of a wild Angolan giraffe. Wild giraffe 1 through 5 were averaged and reported as a single data point along with standard deviation. Heel bulb to distal phalanx volume (HB/P3), Left front lateral claw (LFL), left front medial claw (LFM), right front medial claw (RFM), right front lateral claw (RFL), left rear lateral claw (LRL), left rear medial claw (LFM), right rear medial claw (RRM), right rear lateral claw (RRL)

**Table 2.** Measurements for wild Angolan giraffe and northern giraffe under human care of the volume of the distal phalanx and adipose portion of heel bulb in cubic millimeters. Left front lateral claw (LFL), left front medial claw (LFM), right front medial claw (RFM), right front lateral claw (RFL), left rear lateral claw (LRL), left rear medial claw (LFM), right rear medial claw (RRM), right rear lateral claw (RRL).

| **Specimen** | **Limb** | **P3 volume Cu MM** | **Heel bulb volume Cu MM** |
| --- | --- | --- | --- |
| 2.5 month old | LFL | 23947 | 11500 |
| 2.5 month old | LFM | 24171 | 11058 |
| 2.5 month old | LRL | 20680 | 11425 |
| 2.5 month old | LRM | 20712 | 8948 |
| 2.5 month old | RFL | 24233 | 12046 |
| 2.5 month old | RFM | 25683 | 11375 |
| 2.5 month old | RRL | 20709 | 11259 |
| 2.5 month old | RRM | 21859 | 7754 |
| Namibia Wild Giraffe #1 | LFL | 69553 | 30119 |
| Namibia Wild Giraffe #1 | LFM | 72427 | 28950 |
| Namibia Wild Giraffe #1 | LRL | 53193 | 25353 |
| Namibia Wild Giraffe #1 | LRM | 52411 | 18218 |
| Namibia Wild Giraffe #1 | RFL | 70653 | 32322 |
| Namibia Wild Giraffe #1 | RFM | 72407 | 28180 |
| Namibia Wild Giraffe #1 | RRL | 55438 | 24584 |
| Namibia Wild Giraffe #1 | RRM | 52543 | 18218 |
| Namibia Wild Giraffe #2 | LFL | 71824 | 27553 |
| Namibia Wild Giraffe #2 | LFM | 69648 | 25460 |
| Namibia Wild Giraffe #2 | LRL | 54338 | 25683 |
| Namibia Wild Giraffe #2 | LRM | 51667 | 19226 |
| Namibia Wild Giraffe #2 | RFL | 69836 | 27451 |
| Namibia Wild Giraffe #2 | RFM | 70097 | 25899 |
| Namibia Wild Giraffe #2 | RRL | 52315 | 26123 |
| Namibia Wild Giraffe #2 | RRM | 54064 | 19953 |
| Namibia Wild Giraffe #3 | LFL | 62224 | 24619 |
| Namibia Wild Giraffe #3 | LFM | 62078 | 22951 |
| Namibia Wild Giraffe #3 | LRL | 50974 | 21246 |
| Namibia Wild Giraffe #3 | LRM | 47321 | 15121 |
| Namibia Wild Giraffe #3 | RFL | 64745 | 29652 |
| Namibia Wild Giraffe #3 | RFM | 63964 | 25385 |
| Namibia Wild Giraffe #3 | RRL | 51994 | 22330 |
| Namibia Wild Giraffe #3 | RRM | 48070 | 16368 |
| Namibia Wild Giraffe #4 | LFL | 62755 | 24164 |
| Namibia Wild Giraffe #4 | LFM | 61275 | 23305 |
| Namibia Wild Giraffe #4 | LRL | 50097 | 21233 |
| Namibia Wild Giraffe #4 | LRM | 50141 | 14253 |
| Namibia Wild Giraffe #4 | RFL | 63504 | 25414 |
| Namibia Wild Giraffe #4 | RFM | 61605 | 22417 |
| Namibia Wild Giraffe #4 | RRL | 50795 | 20858 |
| Namibia Wild Giraffe #4 | RRM | 50643 | 20858 |
| Namibia Wild Giraffe #5 | LFL | 55184 | 22495 |
| Namibia Wild Giraffe #5 | LFM | 56052 | 20282 |
| Namibia Wild Giraffe #5 | LRL | 45696 | 19005 |
| Namibia Wild Giraffe #5 | LRM | 43697 | 11989 |
| Namibia Wild Giraffe #5 | RFL | 54309 | 22824 |
| Namibia Wild Giraffe #5 | RFM | 57699 | 19040 |
| Namibia Wild Giraffe #5 | RRL | 45444 | 18402 |
| Namibia Wild Giraffe #5 | RRM | 44757 | 13908 |
| Namibia Wild Giraffe #6 | LFL | 68363 | 20896 |
| Namibia Wild Giraffe #6 | LFM | 69767 | 19320 |
| Namibia Wild Giraffe #6 | LRL | 55103 | 15983 |
| Namibia Wild Giraffe #6 | LRM | 53815 | 10405 |
| Namibia Wild Giraffe #6 | RFL | 68435 | 21408 |
| Namibia Wild Giraffe #6 | RFM | 68371 | 20683 |
| Namibia Wild Giraffe #6 | RRL | 56015 | 14983 |
| Namibia Wild Giraffe #6 | RRM | 54209 | 10214 |

## Discussion

This study aimed to document the anatomy of the lower limb of giraffe and to describe the relative proportions of the hoof capsule and its contents. In doing so we greatly enhanced our understanding of giraffe hoof structures and helped to improve the quality of care provided to giraffe, specifically in terms of hoof health. Our dissections showed some common anatomy between *Giraffa* and other ruminants, specifically in the general anatomy of the tendons and ligaments. However, some of the vascular patterns, particularly the venous drainage of the foot appear to be unique to giraffe. Because giraffe feet are frequently compared to those of bovines and equines when considering hoof care [8], we dissected and sectioned multiple bovine feet for comparison purposes. CT data collected by Gard et al. [15] were also used to compare to CT data collected for this study. The distal limb bones of the giraffe are relatively more gracile than those of the bovine, and also lack the dew claws found in bovines (Fig 39). A prominent feature of the interdigital space in bovines is the large interdigital fat pad (Fig 40a). No appreciable interdigital fat pad is present in giraffe (Fig 40B). The greatest differences, however, are in the morphology of the hoof capsule. The bovine lacks the prominent heel bulb described above in giraffe (Fig 40B). Specifically, there is very little adipose component to the heel in the bovine (Fig 39a and 41A). In contrast, the giraffe not only has a prominent heel bulb but also has a corresponding heel portion on the distal hoof capsule where much of the animal’s weight is supported during locomotion and standing (Figs 39b and 41B). This expanded heel portion of the hoof capsule is a vital weight-bearing component of the sound giraffe foot.

**Figure 39.**
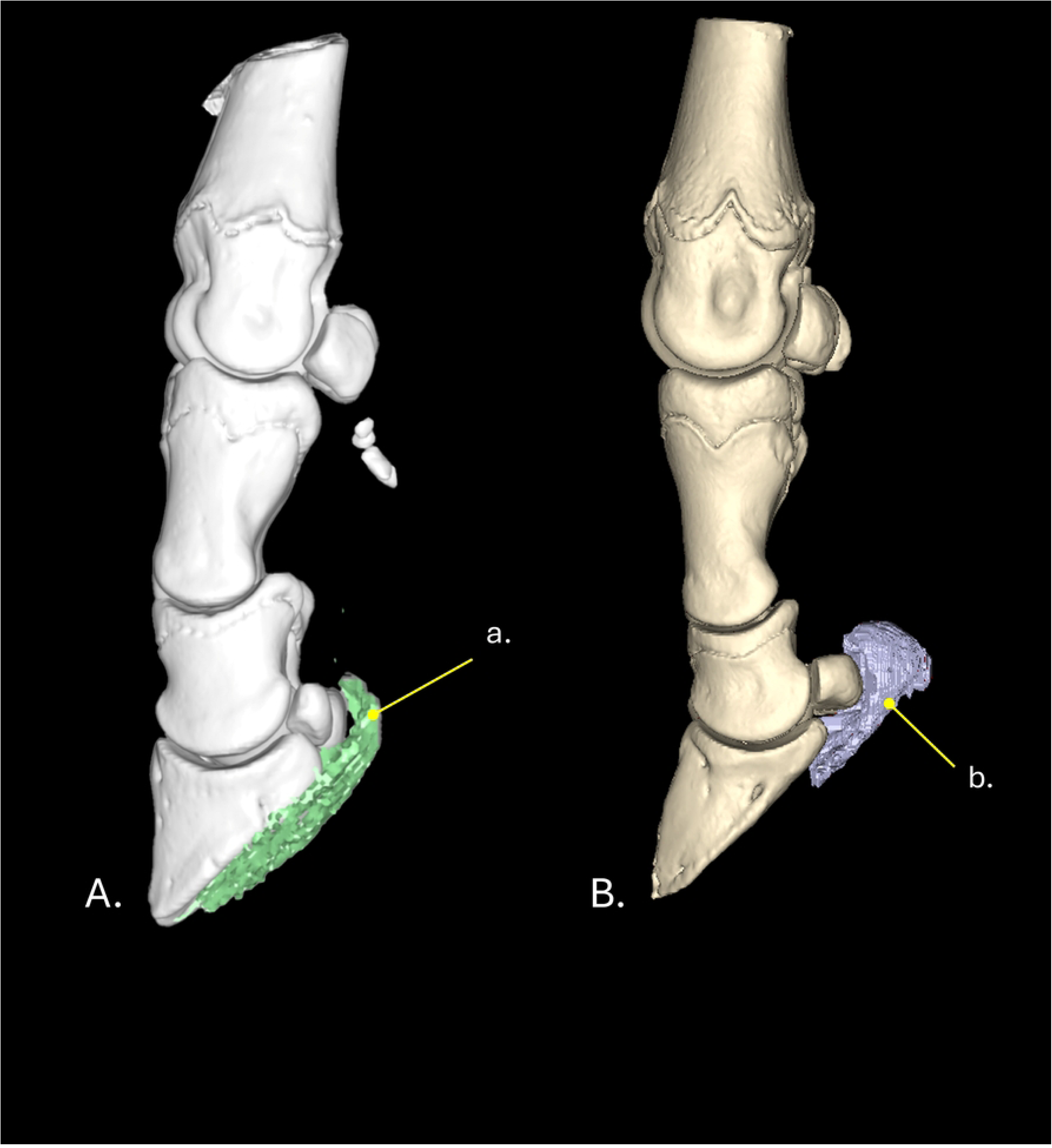
3-D reconstruction of the thoracic limb comparing (A) bovine to (B) wild Angolan giraffe in abaxial view.

**Figure 40.**
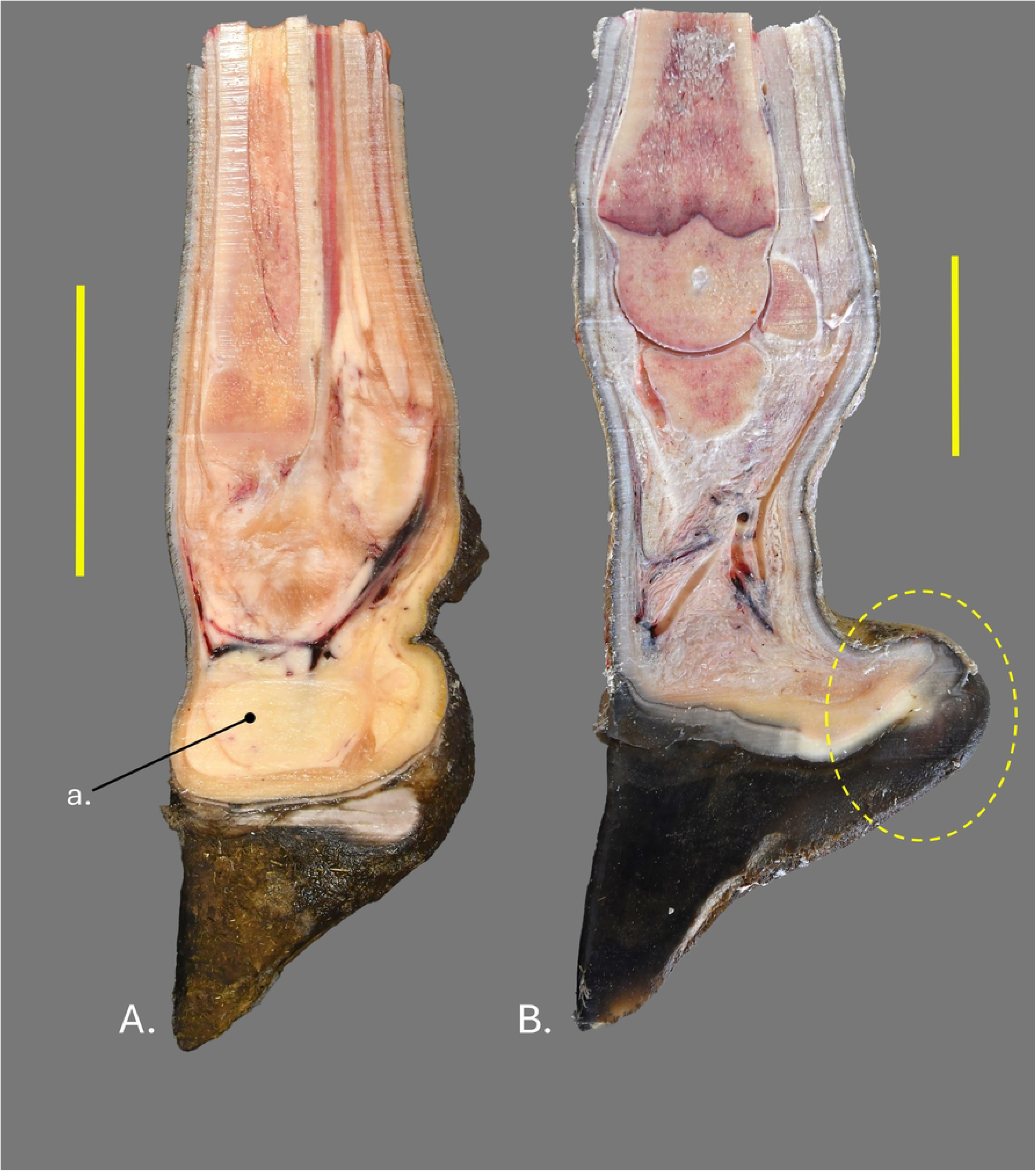
Left thoracic distal limb of a (A) bovine and a (B) wild Angolan giraffe in mid-sagittal view. a- interdigital fat pad. The dotted yellow outline shows the extent of the heel bulb. Scale bar equals 10cm.

**Figure 41.**
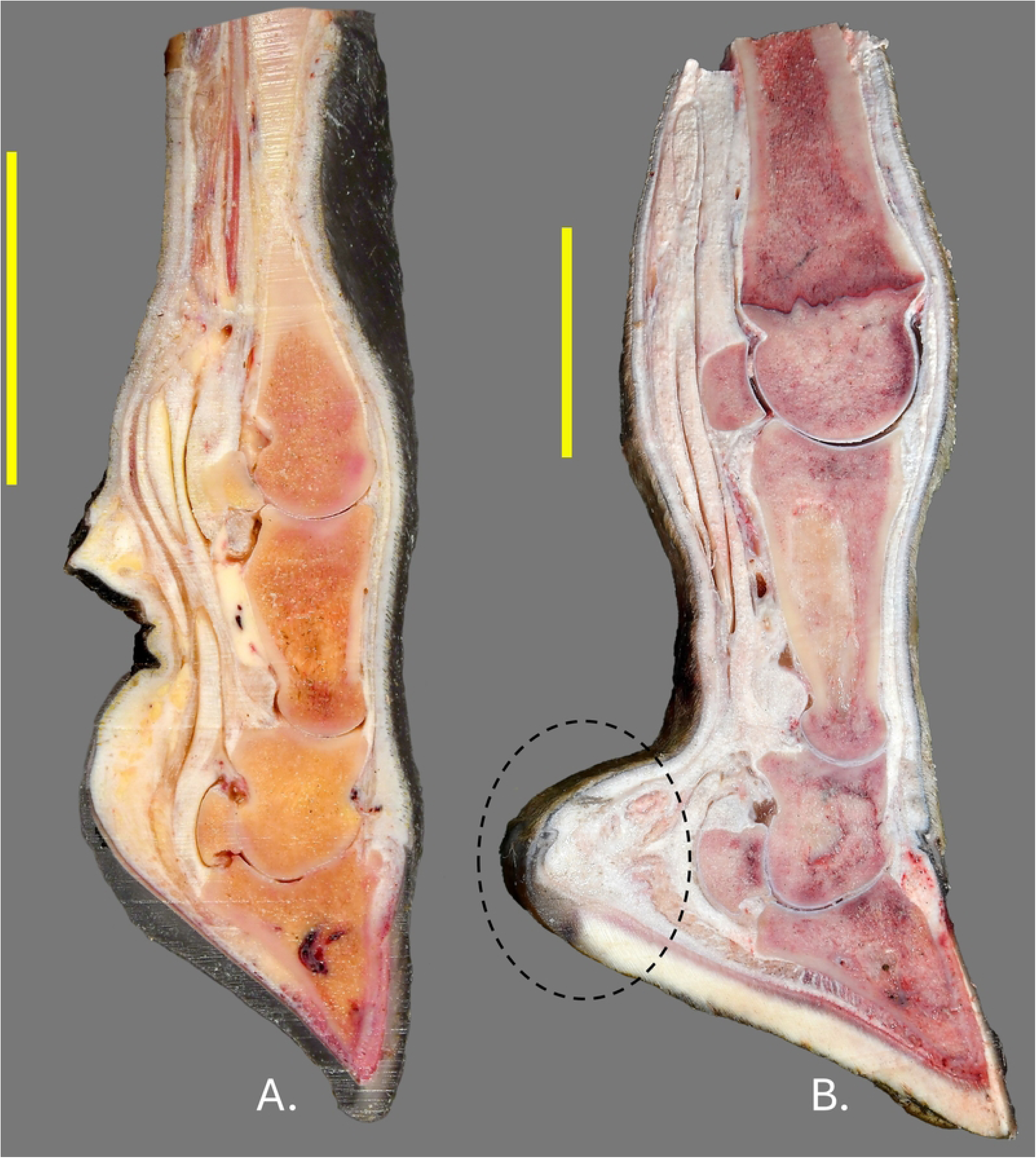
Mid-sagittal section of the medial claw of the left thoracic limb of a (A) bovine and a (B) wild Angolan giraffe. The dotted black outline shows the extent of the heel bulb. Scale bar equals 10cm.

The patterns observed for the relative proportions of the distal phalanges match those of the associated hoof capsules with the lateral hoof capsules either being coequal or larger on both the front and hind feet (Fig 32). We note a distinct difference in the nature of the distal hoof capsule between the concave toe and the softer heel (Fig 32A). We believe this is a significant observation as maintaining a natural transition between the toe and heel could prove vital for giraffe in human care.

Within the hoof capsule we recognize a distinct portion of the digital cushion underlying the distal phalanx that is relatively thin and composed of transverse compartments of adipose tissue separated by dense collagenous connective tissue septa (Figs 33c and 36). Behind this region of the digital cushion, we characterise a portion of the digital cushion contributing to the heel bulb consisting of two large adipose bodies separated by a thick, dense collagenous connective tissue septum and surrounded by a dense connective tissue capsule (Figs. 1a, 2a, 18d, 19d, 33d, 36 and 37a). This interpretation expands on that of Dadone et al. [11] in which the portion of the digital cushion tissue we refer to as part of the heel bulb is considered the complete digital cushion and the thin portion of the digital cushion beneath the distal phalanx is not mentioned. The adipose portion of the heel bulbs of the lateral claws are larger than those of the medial claws in all individuals in this study. Further, as animals increase in age and size, our preliminary data indicates that the relative amount of adipose tissue in the heel bulb decreases. Our hypothesis is that this decrease in relative proportion is due to increased loading of the heel bulb as the animal’s mass increases with age.

## Conclusions

Here we present the first comprehensive published anatomical description of the giraffe distal limb including tendons, ligaments, blood vessels and nerves. This study provides a baseline for future studies, including investigation of pathological distal limbs. Further, we have defined a distinct region of the digital cushion which helps form a significant heel bulb in the giraffe that appears to be unique among large ruminants. We also noted significant differences in the hardness of the sole and heel region of the distal hoof capsule which we believe are related to the mechanical forces experienced by the heel and heel bulb during locomotion.

The biggest strength of this study is the use of normal wild giraffe distal limbs from multiple individuals which provides a solid foundation for comparisons with giraffe in human care. This helps form a baseline of normal anatomy for comparison with human care giraffe distal limbs. Another strength is the use of dissection, CT scanning, 3D modeling and sagittal sections to highlight different aspects of the gross anatomy. CT scanning also allowed for volumetric reconstructions of the expanded digital cushion component of the heel bulbs which we believe will prove valuable in future comparisons with giraffe in human care.

A limitation of the study is the relative young ages of the animals when they were sustainably harvested which could influence our understanding of age related changes to the hoof capsule, sole and heel bulbs. Another limitation is the preponderance of male specimens which did not allow for study of sex related differences.

We believe the heel anatomy is a key indicator of overall hoof health. Future studies will examine older adult human care giraffe both with and without known clinical foot issues to look for changes in the heel bulb region related to pathological processes. If possible, examination of older adult wild giraffe feet would help confirm the trend of decreasing adipose content in the heel bulbs with increased age suggested by this study. Finally, examination of the shape and confirmation of healthy and pathologic adult human care giraffe feet could confirm the importance of the sole to heel hardness transition observed here. This study gives us the baseline by which to compare the feet of giraffe in human care. The ultimate goal of future studies will be guidelines for proper maintenance of human care giraffe feet.

## Acknowledgements

This project was made possible by a joint collaboration between The International Center for the Care and Conservation of Giraffe at Cheyenne Mountain Zoo, the Auburn University College of Veterinary Medicine, and the Giraffe Conservation Foundation. Special thanks to Summer Derrick, Mikey Derrick, Homer Daniels, and Cindy Daniels for their contributions to this project. The Giraffe Conservation Foundation assisted in sourcing the wild specimens, provided space for research and greatly assisted our US research team during their time in Namibia.

Specimens used in the research were kindly provided by Etosha Heights Private Preserve and Okonjati Game Reserve, Namibia under the GCF NCRST permit #RA2024/10/11/02, Certificate #RCIV00042018, and Authorization No #AN2028011402. CT scanning at the Auburn University College of Veterinary Medicine was done by Kim Bryan and CT scanning of wild specimens was done at Medical Imaging Namibia. Deanna Sinclair also helped with dissection and data collection. We also thank the institution who provided the human care specimen used in this study.

